# An EEG Investigation of Neural Dynamics of Empathy Influenced by Congruent and Incongruent Pain Expressions in Autistic and Neurotypical Adults

**DOI:** 10.1101/2025.10.14.681876

**Authors:** Xin Wang, Shelley Xiuli Tong

**Affiliations:** Human Communication, Learning, and Development, Faculty of Education, The University of Hong Kong, Hong Kong, China

**Keywords:** autistic, predictive coding, pain empathy, EEG

## Abstract

Autistic individuals often show difficulties in empathy, but the underlying neural mechanisms of empathy in naturalistic contexts of pain have been less examined. This study employed a kinetic pain empathy paradigm, manipulating the congruence between pain expressions, i.e., body gestures and facial expressions based on a predictive coding framework. We collected EEG data from 51 autistic and 58 neurotypical adults during a pain observation task. Results indicated that autistic and neurotypical adults share a similar neural architecture for empathy processing and conflict resolution, involving an early stage of sensory arousal (i.e., N2 and theta) and a later stage of cognitive reappraisal (i.e., P3). However, the multivariate pattern analysis (MVPA) revealed nuanced but significant between-group differences in neural patterns. Compared to neurotypical peers, autistic adults demonstrated atypical processes in both empathy and conflict resolution. Specifically, they exhibited heightened early emotional arousal but expended greater cognitive effort to evaluate others’ pain. Autistic adults also showed increased alertness to unexpected sensory input and allocated more cognitive resources to resolve prediction errors from incongruent pairings. In contrast, neurotypical adults suppressed unnecessary cognitive efforts for meaningless errors. In summary, autistic adults may experience challenges in efficiently adjusting predictions to the external context, with their neural processing heavily depending on sensory input and less efficient in adapting cognitive resources to evaluate and respond to varied contextual demands.

## Introduction

Autism is characterized as an “empathy disorder” (Gillberg, 1992), but recent research has clarified that the relation between autism and empathy is far more complex (Nicolaidis et al., 2019; Schnitzler & Fuchs, 2024). Additionally, distinct empathy patterns appear in autistic adults and children (Fletcher-Watson & Bird, 2020), but relatively little research on empathy focuses on autistic adults due to challenges in adult diagnosis and a societal focus on children (Brede et al., 2022; Pellicano et al., 2022). Furthermore, behavioural camouflaging in autistic adults (Evans et al., 2024; Khudiakova et al., 2025) may obscure their atypical empathy patterns when using behavioural measurements, which highlights the need for neural-level examinations of empathy processing in autistic adults.

The few neurophysiological research into empathy in autistic adults has largely used isolated and static pain stimuli, such as pictures of body limbs or facial expression (i.e., Liu et al., 2023; Meng et al., 2021), which neglected the complexity of real-life pain experiences. However, two brain studies have suggested that using dynamic and frame-by-frame presentations of two domain pain expressions, i.e., body gestures (moving bodies from standing to bending, Li et al., 2022a) and facial expressions (neutral faces to crying faces, Hadjikhani et al., 2014), can enhance empathy in autistic individuals to comparable levels as neurotypical peers. While autistic individuals struggle with recognizing negative facial expressions (Ghanouni & Zwicker, 2018; Yeung et al., 2020), emerging behavioural research indicates their emotion recognition is largely influenced by the congruence between facial expressions and body gestures (Brewer et al., 2017; Finn et al., 2024). However, no neurophysiological research has examined how the congruence between these two pain expressions influences pain empathy in either neurotypical or autistic adults.

Based on the predictive coding framework, autistic individuals exhibit a strong bias toward local sensory processing and struggle to flexibly evaluate prediction errors between a prediction and the actual input (Marais & Roche-Labarbe, 2025; van Boxtel & Lu, 2013). This can explain their difficulties in understanding the variable and unpredictable experiences and feelings of others (Keysers et al., 2024). However, some well-controlled experiments have shown that autistic individuals have comparable (Ong et al., 2025; Pesthy et al., 2023) or even greater prediction learning abilities (Roser et al., 2015) to neurotypical peers. A key issue with these studies is that they overlooked a higher-order adjustment that individuals weight prediction error based on surrounding social environments (Sapey-Triomphe et al., 2023; Van de Cruys et al., 2014). In social situations, some prediction errors are meaningful and will influence future decisions, while others are insignificant and are likely to be random environmental noise (Kocagoncu et al., 2021). Autistic individuals often struggle to extract these crucial prediction errors (Goris et al., 2018), which may cause difficulties in processing unexpected relations, such as the congruence between multiple pain expressions. There are two domain pain expressions, body gestures and facial expressions, are well recognized (Walsh et al., 2014). Preceding body gestures indicate the resource and location of pain (Rowbotham et al., 2012), while consequent facial expressions convey the affective states of others (Williams, 2002). Although shared empathy networks have been found between these two pain expressions (Jauniaux et al., 2019; Zhou et al., 2020), the influence of their congruence on pain empathy remains underexplored. Therefore, the current study investigates two key questions: (1) To what extent do individuals generate predictions of facial expression based on preceding body gestures, and how do emotional valence and face-body congruence shape the adjustment of prediction error weights? (2) Do autistic individuals demonstrate distinct predictive patterns compared to neurotypical peers, and if so, how? To our knowledge, no neurophysiological studies have examined the predictive processing during pain empathy in either autistic or neurotypical adults. Thus, the current study addressed these gaps by employing time-sensitive EEG measures: event-related potentials (ERPs) and neural oscillations.

ERP research has separated empathy into two subprocesses: an early emotional arousal stage marked by the frontocentral N2 (∼220-350 ms); and late cognitive evaluation stage marked by the parietal P3 (∼400-800 ms), observed in both autistic and neurotypical adults (e.g., Fan et al., 2014; Fan & Han, 2008; Wu et al., 2024). However, the limited research on pain empathy in autistic adults has yielded inconsistent findings, partially due to the wide variety of pain stimuli used. For instance, when presenting static pictures of others’ pain on body limbs, autistic adults showed a larger N2 but a smaller P3 than neurotypical peers (Fan et al., 2014). However, no difference in these ERPs was observed when pain was conveyed through body movements (Li et al., 2022a). Another study using static pictures of others’ pain on faces found a larger P3 in autistic adults than neurotypical peers (Li et al., 2020a), but authors neglected the possibility that the neutral facial expressions might have confounded the perception of an absence of pain. While the parietal P3 has been identified as an indicator of autism severity in adults (Mazer et al., 2024), the absence of P3 effects in some studies could be due to the usage of isolated and static pictures of pain stimuli, which might not be sufficient to engage a full cognitive evaluation.

Additionally, the N2-P3 complex is involved in conflict resolution during facial expression processing. The frontocentral and parietal N2 reflects early conflict detection (∼200-350 ms) while the parietal P3 indicates later cognitive control (∼350-700 ms) (e.g., Brydges et al., 2020; Niessen et al., 2017; Liu et al., 2018). Two ERP studies on neurotypical adults have investigated conflict resolution between facial expressions and body gestures. One observed enhanced N2 amplitudes in incongruent face-body pairings and increased P3 amplitudes in congruent pairings (Li, 2021). Another study observed an enlarged occipital P1 in incongruent pairings (Meeren et al., 2005). However, the use of assembled face-body stimuli in these studies limits their ecological validity. The few ERP studies on autistic adults have only utilized facial expressions in modified Go/Nogo or Flanker paradigms to examine conflict resolution during emotional processing. These studies found impaired prediction processing in autistic adults, evidenced by a prolonged P3 latency in incongruent condition (Fogelson et al., 2019; Thillay et al., 2016). However, No ERP research has yet examined how the N2-P3 complex operates in a dynamic face-body congruence paradigm where both empathy and conflict resolution processes might engage.

Specific neural oscillations, including theta (4-7 Hz) and mu (8-13 Hz), have been identified as neural biomarkers of autism (Parsons et al., 2025; Schwartz et al., 2017). Frontal theta activity, which reflects cognitive control (Cavanagh & Frank, 2014) and emotional sharing in empathy (Mu et al., 2008;), has been shown to co-activate with the N2-P3 complex when processing unpredicted social stimuli (versus predicted ones) and serve as an indicator of the prefrontal cortex maturation in cognitive control (Harper et al., 2014; Wienke et al., 2018). However, no research has examined how the congruence of different pain expressions, which could create predicted and unpredicted pairings, influences theta activity. Additionally, while decreased mu suppression over the sensorimotor cortex has been associated with increased autism severity (Dumas et al., 2014; Wan et al., 2010), a recent meta-analysis identified inconsistencies across these studies, partially due to the usage of non-social stimuli (e.g., hand movements) rather than socially meaningful stimuli (e.g., dynamic facial expressions; Lockhart et al., 2024). For instance, one study on pain empathy using static pictures of injured body limbs found no differences in mu suppression between autistic and neurotypical adults (Fan et al., 2014), suggesting that static social stimuli may not be sufficient to activate autism-related sensorimotor resonance during empathy processing. While ERP studies dominate research on pain empathy and conflict resolution, frequency-based EEG investigations remain scarce, especially in autistic adults. By examining both temporal (e.g., N2 and P3) and oscillatory (e.g., theta and mu) neural dynamics, the present study provides a more comprehensive understanding of how autistic and neurotypical adults process conflict resolution and empathy during the observation of congruent and incongruent pain expressions.

This study advanced the understanding of the neural mechanisms underlying pain empathy in both autistic and neurotypical adults through two key innovations in experimental paradigm and analytical approach. Specifically, we developed dynamic pain animations that integrated naturalistic pain scenarios with congruent or incongruent pain expressions between preceding body gestures and subsequent facial expressions. We also integrated multivariate pattern analysis (MVPA) with traditional univariate analyses, including both event-related potentials (ERPs) and event-related spectral perturbations (ERSPs). Two research questions are raised: (1) How does the congruence between body gestures and facial expressions with pain influence the neural dynamics of pain empathy? (2) To what extent do autistic and neurotypical adults differ in their neural responses to congruent and incongruent pain expressions? We hypothesized that there would be a significant interaction between congruence and pain, activating neural mechanisms for both pain empathy and conflict resolution, as characterized by the N2-P3 complex, theta, and mu activations. Additionally, we hypothesized that autistic adults would show distinct neural patterns compared to neurotypical peers, manifesting as weaker empathy and impaired conflict resolution.

## Method

### Participants

Fifty-four autistic adults (26 males; *M* = 26.03 years, *SD* = 7.82 years) and sixty neurotypical adults (31 males; *M* = 26.03 years, *SD* = 7.82 years) participated in the present study. An adult was included in the autistic group if he/she (1) received a medical diagnose of autism or autism spectrum disorder (ASD) by qualified professionals (e.g., a psychologist or psychiatrist); and (2) exhibited ASD symptoms via the Autism Diagnostic Observation Schedule (2nd edition, ADOS-2; Lord et al., 2012), administered by certificated researchers within one week prior to EEG data collection. An adult was included in the neurotypical group if he/she (1) had no clinical diagnosis or suspicion of ASD, (2) was absence of psychiatric disorders or neurodevelopmental disorders, and (3) showed no ASD symptoms based on the ADOS-2 screening. All participants met the following criteria: (1) nonverbal IQ was greater than 90 measured by the 30-item matrix reasoning subset of the Wechsler Abbreviated Scale of Intelligence (WASI-II, Wechsler, 2011); (2) had normal or corrected-to-normal vision; (3) had no history of brain injury or surgery; (4) took no psychoactive medications in the last one week. However, due to the high prevalence of specific comorbidities (i.e., anxiety, depression, and ADHD) in autistic adults (Pehlivanidis et al., 2020; Suen et al., 2024), autistic participants with histories of these commodities but reported no active symptoms during the assessment were included, i.e., two with anxiety, one with depression, and six with ADHD during early childhood. Written informed consent was obtained from all participants, who were compensated with $ 20 USD for their engagement. This study received ethical approval from the Human Research Ethics Committee of the authors’ university.

Due to excessive artifacts (e.g., eye blinks and muscle contractions) or task incompletion, three autistic adults and two neurotypical adults were removed as fewer than 60% of trials retained after EEG preprocessing. The final analysis consisted of fifty-one autistic adults (24 males; *M* = 25.57 years, *SD* = 7. 20 years) and fifty-eight neurotypical adults (29 males; *M* = 25.27 years, *SD* = 5.44 years). Autistic adults obtained significantly higher ADOS scores in communication (*M* = 2.39, *SD* = 1.18, *t* = 14.40, *p* < .001, *d* = 2.76) and reciprocal social interaction (*M* = 5.84, *SD* = 1.84, *t* = 23.60, *p* < .001, *d* = 4.52) compared to neurotypical adults (communication: *M* = 0.09, *SD* = 0.28; reciprocal social interaction: *M* = 0.07, *SD* = 0.32), indicating significant diagnostic differences between two groups.

### Experimental Design and Evaluation

Each animation lasts for 9 seconds (s) with three phases, including the first antecedent cause phase (0-3 s) setting a neutral scene, the second body gesture phase (3-6 s) depicting a pain or no-pain event followed by corresponding body gestures (BG), and the final facial expression phase (6-9 s) depicting the protagonist’s facial expressions (FE) with pain or with no-pain. Four experimental conditions were developed based on the congruence between the last two phases. Therefore, we have (1) BG-FE-condition with congruent body gestures in response to no-pain events followed by facial expressions with no-pain, (2) BG-FE+ condition with incongruent body gestures in response to no-pain events followed by facial expressions with pain, (3) BG+FE-condition with incongruent body gestures in response to pain events followed by facial expressions with no-pain, and (4) BG+FE+ condition with congruent body gestures in response to pain events followed by facial expressions with pain.

Thus, seventy-six animated videos (i.e., 19 stimuli sets × 4 conditions = 76 stimuli) depicted four experimental conditions, featuring different protagonists experiencing various pain events in a variety of real-life settings, such as homes, parks, and schools.

### Stimuli Evaluation

A total of 40 neurotypical university students (20 males; *M* = 24.87 years, *SD* = 3.64 years) participated in the online evaluation animation comprehensibility and applicability.

We selected 19 of 34 animation sets with BG-FE-condition being perceived as *No-pain* from all participants when rating “*Did the protagonist experience pain? (Yes-No).*” The 76 animations (19 stimuli sets × 4 conditions = 76 stimuli) were rated with above-average comprehensibility scores (question 2: *How well did you understand the animation? M* = 3.25 > 2, *SD* = 0.96, *t* = 76.63, *p* < .001). Four conditions demonstrated progressively higher degree of pain and imagined pain from BG-FE-to BG-FE+ to BG+FE-till BG+FE+ for following questions “*How much pain do you think the protagonist experienced?*” and “*How well would you handle the pain if you were that protagonist?*”(Q3: |β|s > 0.66, *Se*s = 0.15, |*z*|s > 44.28, *p*s < .001; Q4: |β|s > 0.43, *SE*s = 0.12, |*z*|s > 3.51, *p*s < .003). The protagonists’ emotional states (“*How was the emotional state experienced by the protagonist?*”) under conditions BG-FE+ and BG+FE+ were significantly more unpleasant than those under conditions BG-FE- and BG+FE-(|β|s > 0.84, *SE*s = 0.11, |*z*|s > 7.68, *p*s < .001). In sum, the 19 animation sets remained as formal stimuli.

### Experimental Procedure

All participants completed a pain empathy task on a high-resolution computer (1,600 × 1,200 dpi) in an electromagnetic interference acoustic booth. The task consisted of passive video viewing interspersed with active rating periods.

Specifically, each animated video (9,000 ms) was passively presented, preceded by an approximately 1,500-3,500 ms display of fixation “+” and a blank screen (1,000 ms-2,000 ms), and followed by another blank screen (500 ms-1,000 ms). The longer duration of blank screen preceding video viewing allowed sufficient time for brain activation to return to baseline.

After viewing each video, participants were required to rate four questions, which assessed a hierarchical empathy processing, from sensory perception (“*How much does the protagonist suffer?*”), affective recognition (“*What is the emotion of the protagonist?*”), self-other distinct (“*If you are the protagonist, how much do you suffer?*”), to empathic concern (“*How sorry are you for the protagonist?*”).

All 76 animated videos were played twice (i.e., 152 trials) in a pseudo-randomized order without consecutive repetition of the same video, ensuring adequate condition-wise trial numbers and minimizing video-specific variations. Before the formal task, four extra practice trials (one per condition) oriented participants to the experiment procedure.

### EEG Data Acquisition and Preprocessing

EEG data were recorded using a NeuroScan system with the 64-electrode Ag/AgCl EEG caps following the 10-20 system. The online sampling rate was 1000 Hz, and the online reference was on the vertex electrode. The impedance of all channels was adjusted under 10 kꭥ prior to the formal recording.

Figure 1 shows the EEG data preprocessing protocol with identical protocols for both autistic and neurotypical groups, except for modifications of the band-pass filter thresholds. Specifically, given excessive neural noise in autistic individuals (Raul et al., 2024), we employed a 0.1-60 Hz band-pass filter which was applied in prior EEG studies in autism (Djemal et al., 2017; Gabard-Durnam et al., 2019). The number of rejected ICs for autistic adults (*M* = 4.41, *SD* = 2.15, *Min* = 1, *Max* = 12, *t* = 0.63, *p* = .735) was not significantly different from the one for neurotypical adults (*M* = 4.67, *SD* = 2.17, *Min* = 1, *Max* = 11). There was no significant difference of remaining trials between autistic (*M* = 145, *SD* = 12.90) and neurotypical groups (*M* = 150, *SD* = 9.81, *t* = −0.39, *p* = .701).

**Figure 1.**
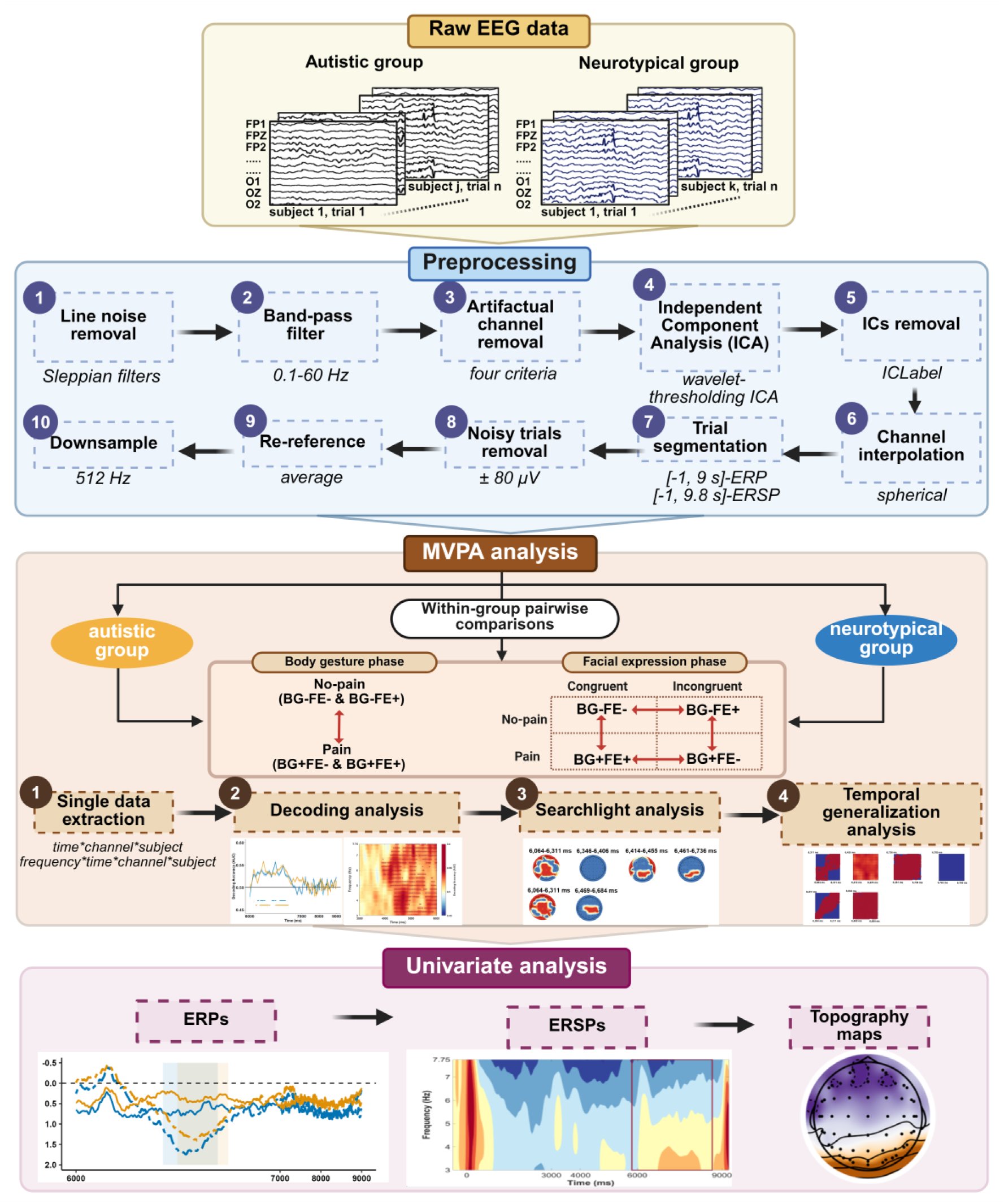
EEG Data Analysis Pipeline.

### Multivariate Pattern Analysis (MVPA)

To examine group-specific temporospatial neural representations underlying different experimental conditions, we performed condition-wise MVPA for autistic and neurotypical groups independently using the MVPA-light toolbox (Treder, 2020) in MATLAB. Specifically, the predefined condition pairs (Figure 1) included one contrast between pain and no-pain events during the body gesture phases (BG+FE- and BG+FE+ vs. BG-FE- and BG-FE+), and four contrasts during the facial expression phase: (1) a comparison between congruence and incongruence under no-pain events (BG-FE-vs. BG-FE+), (2) a comparison between congruence and incongruence under pain events (BG+FE+ vs. BE+FE-), (3) a comparison between two congruent conditions (BG+FE+ vs. BG-FE-), and (4) a comparison between two incongruent conditions (BG-FE+ vs. BG+FE-).

To investigate neural patterns from both temporal and frequency dimensions, we conducted MVPA separately on temporal-based and frequence-based EEG data. While temporal-based EEG data were derived directly from epoch segmentations, frequency-based data were computed through time-frequency decomposition using *newtimef()* function in MATLAB. The decomposition employed Morlet wavelets with five cycles ([3 0.8], Kronland-Martinet et al., 1987) on the 3–30 Hz. Finally, we only extracted post-stimulus single-trial EEG data (i.e., 0-9,000 ms) from either time-domain or time-frequency domain across three frequency bands, i.e., theta (3-8 Hz), mu (8-12 Hz), and beta (13-30 Hz).

A comprehensive MVPA consisted of three sequential steps: decoding analysis, searchlight analysis and temporal generalization analysis (Figure 1). A linear discriminant analysis (LDA) classifier was used for classification among all three steps. First, we performed “undersampling” process to randomly assign equal numbers of trials to training and testing subsets. Next, ten times k-fold cross-validation (k = 5) were applied. The area under the ROC curve (AUC) represented the classification performance. All time-series AUC values were compared against chance-level classification (AUC = 0.5) using non-parametric counterpart Wilcoxon cluster permutation tests at both individual and group levels, with the significance threshold at *p* < .05 and the permutation at 2,000 iterations.

The time-resolved decoding analysis was conducted at each time point for time-domain data and was performed at each time-frequency point within three frequency bands (i.e., theta, mu, and beta) separately for time-frequency data. After identifying time periods with significant classification performances, the searchlight analysis was performed for each EEG channel during these specific periods separately. Finally, temporal generalization analysis was employed for periods on these channels with significant classification performances. Only periods with acceptable classification stability, i.e., significant classifications along and below the diagonal line on the generalized matrix, were used during subsequent analyses.

### Univariate Analysis

To characterize neural mechanisms associated with significant classification results, we performed univariate analyses of event-related potentials (ERPs) and event-related spectral perturbations (ERSPs) at the identified time periods and channels with significant classification performance. ERP data was baseline-corrected according to pre-stimulus periods (i.e., −500 ms-0 ms) and only standardized ERP waveforms were analysed. ERSP data was baseline-corrected according to pre-stimulus periods (i.e., −500 ms-100 ms) and transferred to decibel units via the formulation: [dB power = 10*log10 (power/baseline)]. Then, condition comparison employed linear mixed models (LMMs) (Baayen et al., 2008) via the lmerTest package (Kuznetsova et al., 2017) in R. Fixed effects were the specific condition contrasts and, where applicable, group (autistic vs. neurotypical). Gender and age were included as covariates in the models. Random effects accounted for individual differences. Dependent variables were either time window-averaged ERP amplitudes or time window-averaged ERSP power. The *anova*() function in R examined fixed effects by performing Type III F-tests with either Satterthwaite or Kenward-Roger adjustments for degrees of freedom (Kuznetsova et al., 2017). Post-hoc comparisons used the emmeans package (Lenth, 2023), with Tukey’s HSD adjustment to control for multiple comparisons. The *confint*() function computed 95% confidence intervals.

For behavioural ratings, LMMs analysed ratings across four questions, with condition (four levels) and group (autistic vs. neurotypical) as the fixed effects and participant as the random effect, following the same statistical approach as described above.

## Results

### Behavioural Results

Autistic adults rated significantly less pain for protagonists under the conditions BG+FE-(*M* = 2.32, *SD* = 0.97, β = −0.20, *SE* = 0.08, *z* = −2.48, *p* = .013, 95% CI = [-0.35, −0.04]) and BG+FE+ (*M* = 3.17, *SD* = 1.10; β = −0.21, *SE* = 0.08, *z* = −2.58, *p* = .010, 95% CI = [-0.36, −0.05]) than neurotypical adults (BG+FE-: *M* = 2.52, *SD* = 1.00; BG+FE+: *M* = 3.38, *SD* = 1.04). Similarly, autistic adults rated significantly less pain for themselves when imagining they were protagonists under the conditions BG+FE-(*M* = 2.47, *SD* = 1.12, β = −0.20, *SE* = 0.09, *z* = −2.32, *p* = .020, 95% CI = [-0.37, −0.03]) and BG+FE+ (*M* = 3.00, *SD* = 1.19, β = −0.21, *SE* = 0.09, *z* = −2.44, *p* = .015, 95% CI = [-0.39, −0.04]) than neurotypical adults (BG+FE-: *M* = 2.69, *SD* = 1.11; BG+FE+: *M* = 3.24, *SD* = 1.13). Autistic adults rated significantly less sad for protagonists under the conditions BG+FE+ (*M* = 1.96, *SD* = 0.87, β = 0.26, *SE* = 0.05, *z* = 4.88, *p* < .001, 95% CI = [0.03, 0.23]) and BG-FE+ (*M* = 1.87, *SD* = 0.88, β = 0.35, *SE* = 0.05, *z* = 6.53, *p* < .001, 95% CI = [0.15, 0.36]) than neurotypical adults (BG+FE+: *M* = 1.68, *SD* = 0.71; BG-FE+: *M* = 1.67, *SD* = 0.66). Autistic adults exhibited significant less empathic concern to protagonists under the conditions BG+FE-(*M* = 1.75, *SD* = 0.94, β = −0.40, *SE* = 0.07, *z* = −6.17, *p* < .001, 95% CI = [-0.53, −0.27]) and BG+FE+ (*M* = 2.16, *SD* = 1.19, β = −0.58, *SE* = 0.07, *z* = −8.89, *p* < .001, 95% CI = [-0.71, −0.45]) than neurotypical adults (BG+FE-: *M* = 1.82, *SD* = 1.06; BG+FE+: *M* = 2.70, *SD* = 1.25).

### Multivariate and Univariate Analyses of Temporal-based EEG Data

Results of temporal-based MVPA and univariate analysis were reported according to different phases and condition contrasts.

### Body Gesture Phase

Autistic adults demonstrated a significant above-channel classification in distinguishing body gestures in response to pain events versus no-pain events between 5,348 ms and 5,389 ms (*p* clusters < .05, *M*ean AUC = 0.52, Figure 2a), particularly in the right frontocentral region (*p* clusters < .05, *M*ean AUC = 0.52, Figure 2b). However, no further univariable analysis was conducted as the significance did not persist across time during the period (Figure 2c).

**Figure 2.**
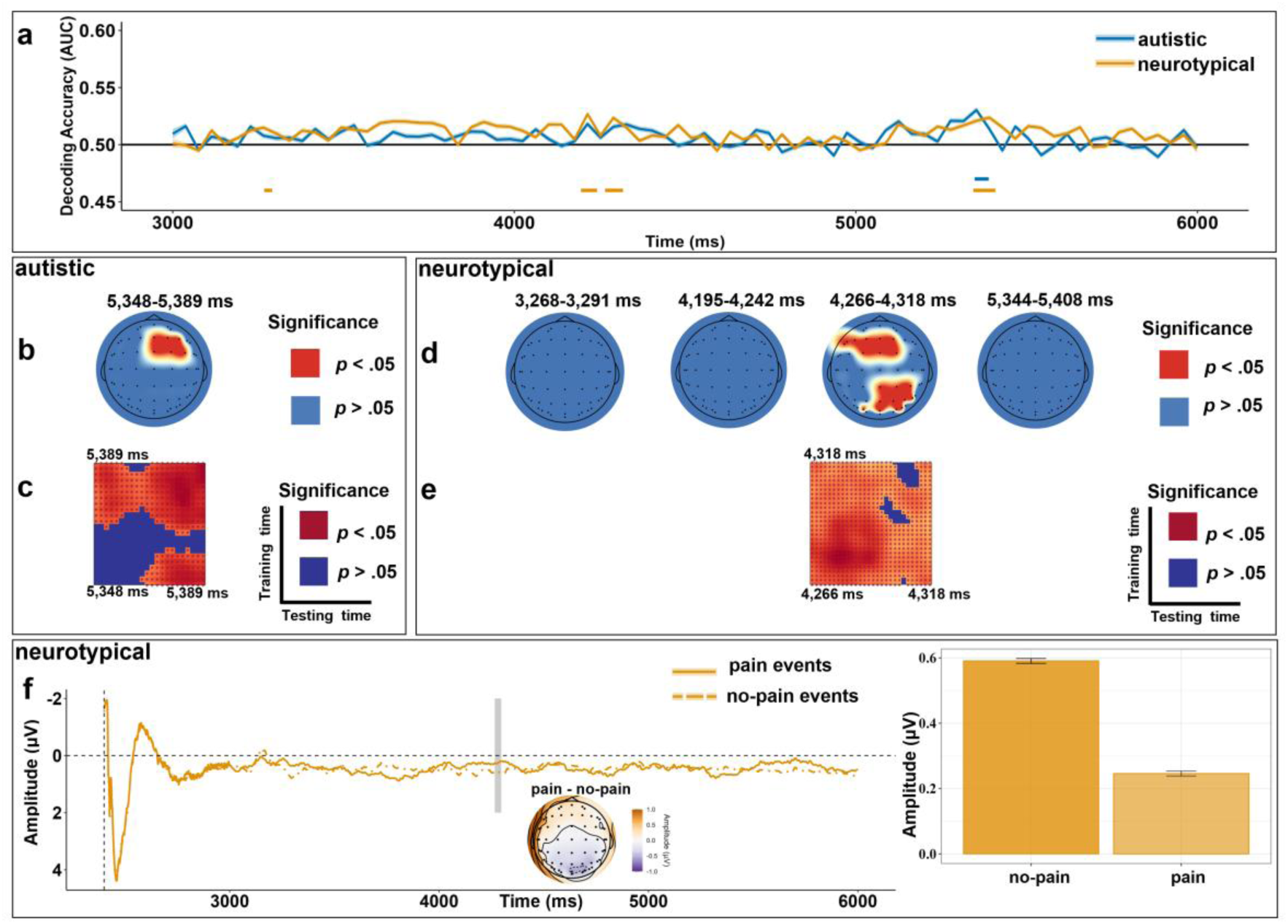
Temporal-based MVPA and ERP Results During the Body Gesture Phase. *Note*. Panel a: Classification accuracy for pain vs. no-pain events in autistic (blue lines) and neurotypical (yellow lines) groups, where significant time clusters (*p* clusters < .05) are marked with horizontal bars below the decoding curves. Searchlight results for autistic (Panel b) and neurotypical (Panel d) groups reveal channels with (red dots, *p* clusters < .05) and without significant (blue dots, *p* clusters > .05) contributions during certain periods. Temporal generalization matrices for autistic (Panel c) and neurotypical (Panel e) groups mark periods with significant classification stability (*p* clusters < .05) in dark red, while unstable periods (*p* clusters > .05) in dark blue. Panel f: ERP waveform displays neural responses to pain and no-pain events over time recorded at parieto-occipital channels for neurotypical adults (averaged value in the right bar chart). The topographic map illustrates the amplitude distribution from 4,266 ms to 4,297 ms (highlighted by a grey bar in the ERP waveform).

Neurotypical adults exhibit significant classification (*M*ean AUC = 0.52, Figure 2a) with channel-specific contributions in the frontocentral and parieto-occipital regions (*M*ean AUC = 0.52, Figure 2d), sustaining from 4,266 ms to 4,297 ms (*p* clusters < .05, Figure 2e). We observed the peak of one negative ERP merged between 4,266 ms and 4,297 ms at channels PZ, P2, P4, P6, P8, PO5, PO3, POZ, PO4, and PO6 (Figure 2f), with a significantly greater negative deflection for pain (*M* = 0.24, *SD* = 0.46, β = 0.34, *SE* = 0.09, *z* = 3.74, *p* < .001, 95% CI = [0.16, 0.53]) than no-pain events (*M* = 0.59, *SD* = 0.55).

### Facial Expression Phase

#### Comparing congruence between two pain expressions under no-pain events (BG-FE-vs. BG-FE+) between two groups

Autistic adults exhibited significant above-channel classification in distinguishing incongruent facial expressions with pain (BG-FE+) versus congruent ones with no-pain (BG-FE-) during four periods with sustainable generalization across time (*p* clusters < .05, Figure 3a, c) and channel-specific contributions in the centroparietal region (*p* clusters < .05, *M*ean AUC = 0.53, Figure 3b): (1) 6,146 ms-6,180 ms (*M*ean AUC = 0.53), (2) 6,425 ms-6,471 ms (*M*ean AUC = 0.54), (3) 6,480 ms-6,510 ms (*M*ean AUC = 0.54), and (4) 6,576 ms-6,695 ms (*M*ean AUC = 0.54).

**Figure 3.**
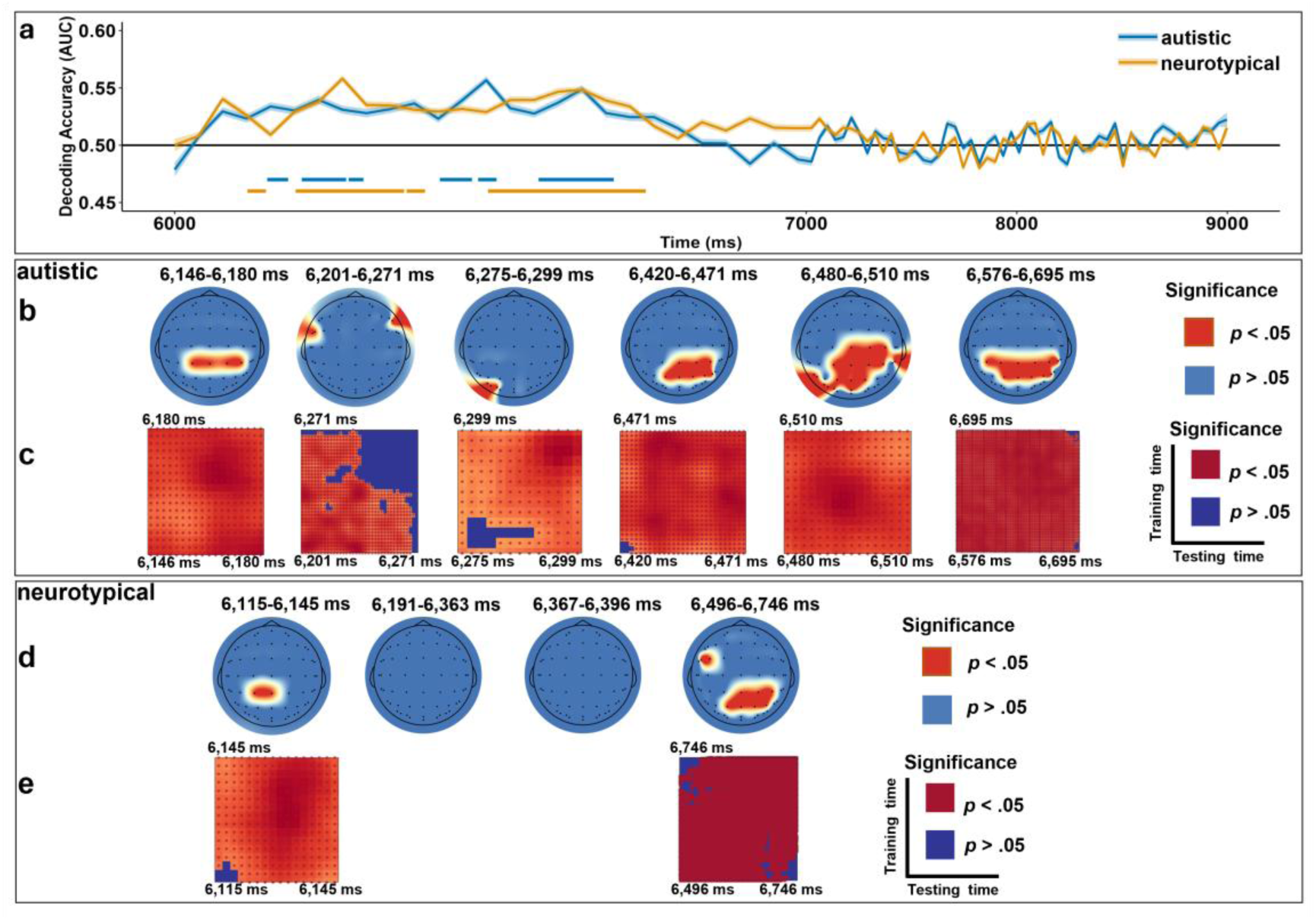
Temporal-based MVPA Results Comparing BG-FE- and BG-FE+ During the Facial Expression Phase. *Note*. Panel a: Classification accuracy in autistic (blue lines) and neurotypical (yellow lines) groups, where significant time clusters (*p* clusters < .05) are marked with horizontal bars below the decoding curves. Searchlight results for autistic (Panel b) and neurotypical (Panel d) groups reveal channels with (red dots, *p* clusters < .05) and without significant (blue dots, *p* clusters > .05) contributions during certain periods. Temporal generalization matrices for autistic (Panel c) and neurotypical (Panel e) groups mark periods with significant classification stability (*p* clusters < .05) in dark red, while unstable periods (*p* clusters > .05) in dark blue. The x-axis employs a piecewise linear transformation where the interval 7,000-9,000 ms is compressed for a better visualization.

Neurotypical adults exhibited significant classification during two periods with sustainable generalization across time (*p* clusters < .05, Figure 3a, e) and channel-specific contributions (*p* clusters < .05, Figure 3d): (1) 6,119 ms-6,145 ms (*M*ean AUC = 0.52) in the centroparietal region (*M*ean AUC = 0.53), and (2) 6,496 ms-6,746 ms (*M*ean AUC = 0.53) in the centroparietal and left frontotemporal regions (*M*ean AUC = 0.54).

During the first period: 6,146 ms to 6,180 ms for autistic adults, and 6,119 ms-6,145 ms for neurotypical adults, we observed the peak of one negative ERP at channels CP3, CP1, CPZ, CP2, and CP4 (Figure 4). Results showed that the incongruent condition (BG-FE+; *M* = −0.69, *SD* = 0.75, β = −0.64, *SE* = 0.08, *t* = - 8.20, *p* < .001, 95%CI = [-0.79, −0.48]) elicited significantly larger negative deflection than the congruent condition (BG-FE-; *M* = −0.06, *SD* = 0.62). No significant group-relevant main or interaction effect was found (*p*s > .05).

**Figure 4.**
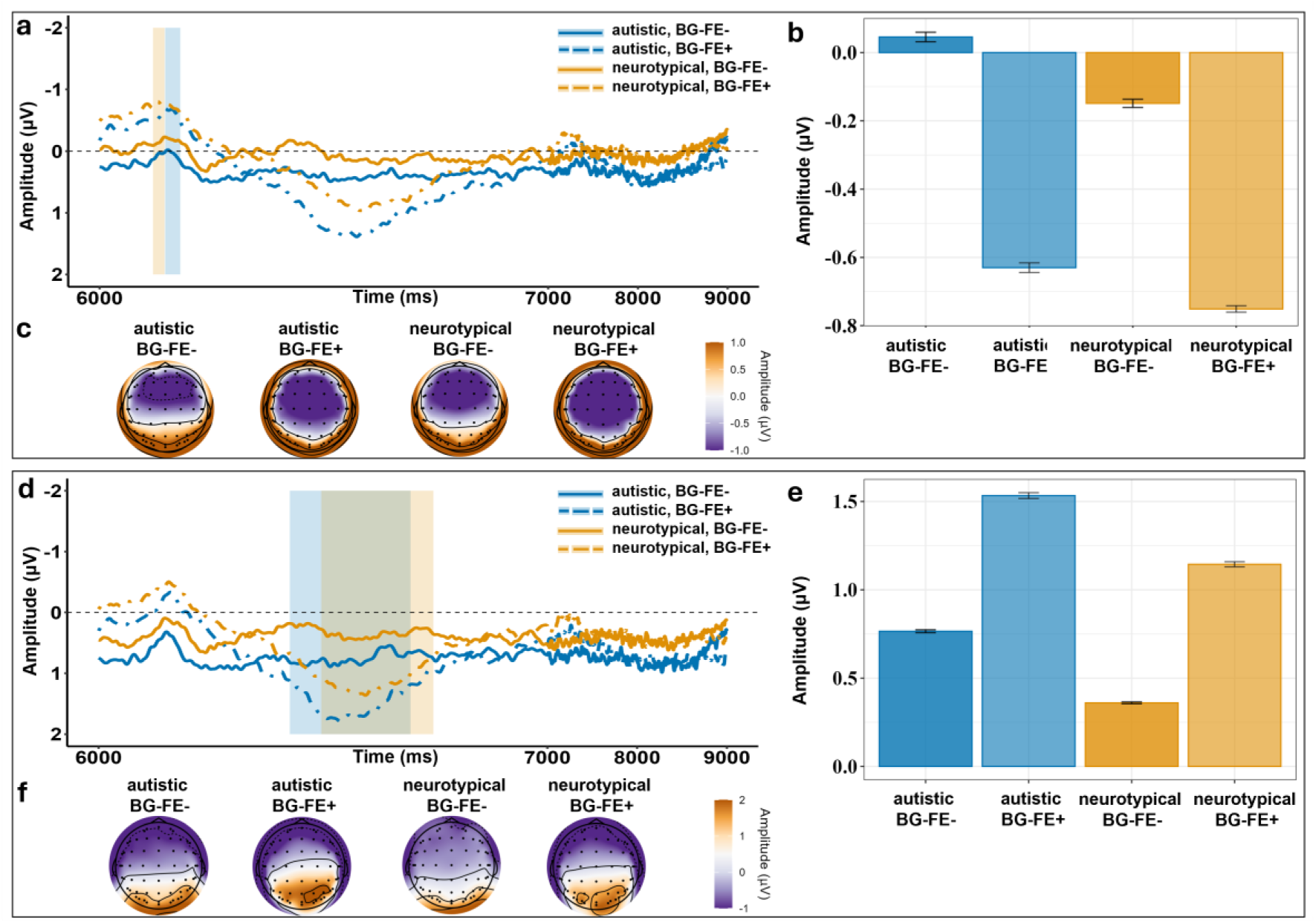
ERP Results Comparing BG-FE- and BG-FE+ During the Facial Expression Phase. *Note*. ERP waveforms display neural responses across two conditions and two groups over time, at the centroparietal channels (Panel a), and at the centroparietal and parietal channels (Panel d). The topographic maps (Panel c) illustrate the amplitude distribution from 6,146 ms to 6,180 ms for autistic adults (highlighted by the light blue bar in Panel a) and from 6,119 ms to 6,145 ms for neurotypical adults (highlighted by the light yellow bar in Panel a). The topographic maps (Panel f) illustrate the amplitude distribution from 6,426 ms to 6,695 ms for autistic adults (highlighted by the light blue bar in Panel d) and from 6,496 ms to 6,746 ms for neurotypical adults (highlighted by the light yellow bar in Panel d). The bar charts (Panels b and e) compare neural responses between two conditions and two groups during corresponding time windows and at selected channels. The x-axis employs a piecewise linear transformation where the interval 7,000-9,000 ms is compressed for a better visualization.

Additionally, we observed the peak of one positive ERP at channels CP3, CP1, CPZ, CP2, CP4, P3, P1, PZ, P2, and P4 in both groups: 6,426 ms to 6,695 ms for autistic adults and 6,496 ms to 6,746 ms for neurotypical adults (Figure 4). Autistic adults (*M* = 1.15, *SD* = 0.79, β = 0.40, *SE* = 0.11, *t* = 3.74, *p* < .001, 95%CI = [0.19, 0.61]) exhibited a significantly larger positive deflection than neurotypical adults (*M* = 0.75, *SD* = 0.68). The incongruent condition (BG-FE+; *M* = 1.33, *SD* = 0.70, β = 0.78, *SE* = 0.06, *t* = 13.66, *p* < .001, 95%CI = [0.66, 0.89]) elicited a significantly larger positive deflection than the congruent condition (BG-FE-; *M* = 0.55, *SD* = 0.60). No significant interaction effect was found (*p* > .05).

#### Comparing congruence between two pain expressions under pain events (BG+FE+ vs. BG+FE-) between two groups

Autistic adults exhibited significant above-channel classification in distinguishing congruent facial expressions with pain (BG+FE+) versus incongruent ones with no-pain (BG+FE-), which significantly sustained across time during three periods with channel-specific contributions (*p* clusters < .05, Figure 5a-c): (1) 6,010 ms-6,207 ms (*M*ean AUC = 0.55) in the centroparietal region (*M*ean AUC = 0.54), (2) 6,213 ms-6,617 ms (*M*ean AUC = 0.56) from the frontal to occipital regions (*M*ean AUC = 0.54), and (3) 6,621 ms-6,664 ms (*M*ean AUC = 0.54) in the anterior and parietal regions (*M*ean AUC = 0.53).

**Figure 5.**
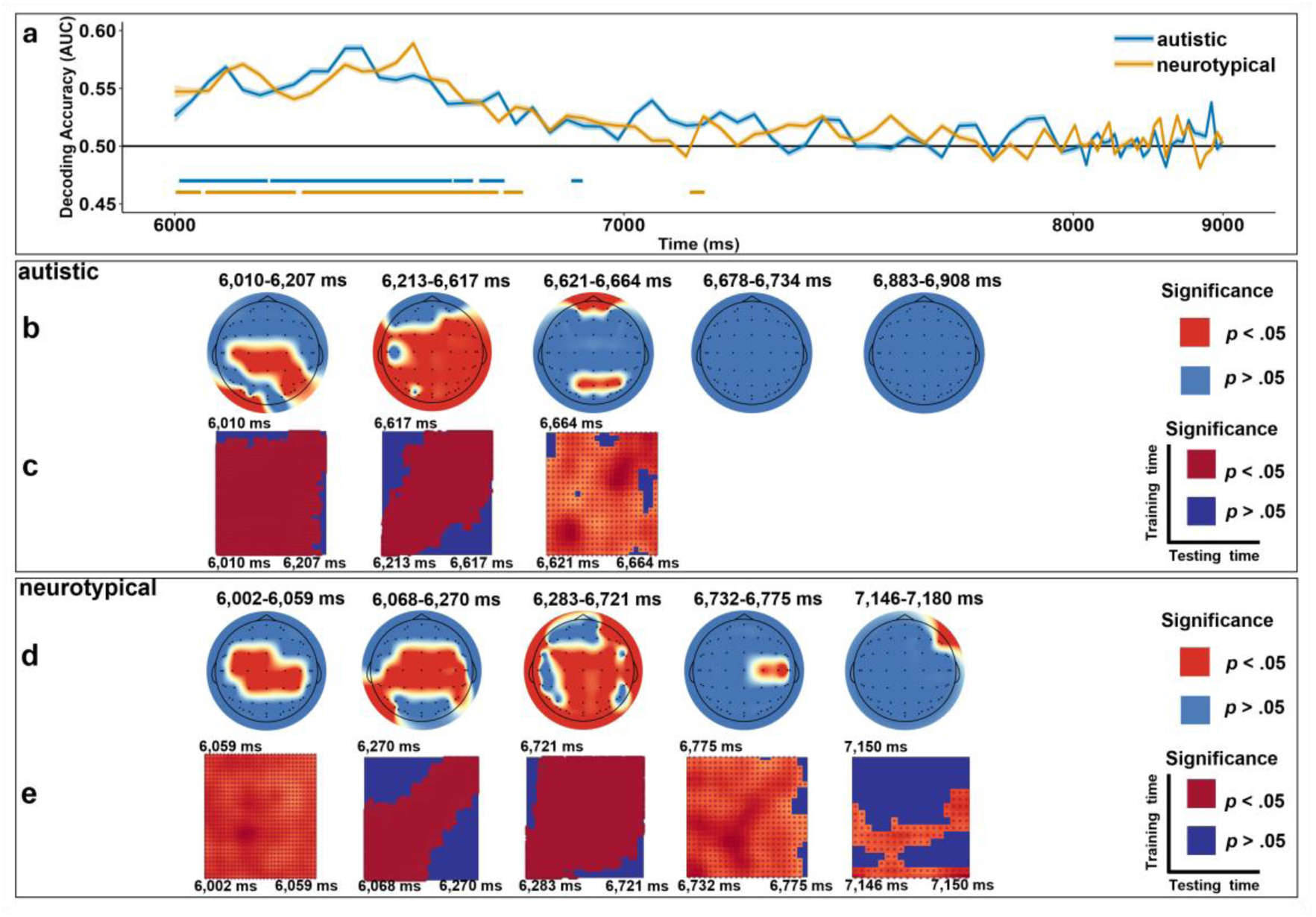
Temporal-based MVPA Results Comparing BG+FE+ and BG+FE-During the Facial Expression Phase. *Note*. Panel a: Classification accuracy in autistic (blue lines) and neurotypical (yellow lines) groups, where significant time clusters (*p* clusters < .05) are marked with horizontal bars below the decoding curves. Searchlight results for autistic (Panel b) and neurotypical (Panel d) groups reveal channels with significant (red dots, *p* clusters < .05) and without significant (blue dots, *p* clusters > .05) contributions during certain periods. Temporal generalization matrices for autistic (Panel c) and neurotypical (Panel e) groups mark periods with significant classification stability (*p* clusters < .05) in dark red, while unstable periods (*p* clusters > .05) in dark blue. The x-axis employs a piecewise linear transformation where the interval 8,000-9,000 ms is compressed for a better visualization.

Neurotypical adults showed significant classification during four periods, which significantly sustained across time with channel-specific contributions (*p* clusters < .05, Figure 5a, d, e): (1) 6,002 ms-6,059 ms (*M*ean AUC = 0.54) in the frontocentral region (*M*ean AUC = 0.54), (2) 6,068 ms-6,270 ms (*M*ean AUC = 0.55) from the frontal to parietal regions (*M*ean AUC = 0.54), (3) 6,283 ms-6,721 ms (*M*ean AUC = 0.56) in the most frontal to occipital regions (*M*ean AUC = 0.54), and (4) 6,732 ms-6,775 ms (*M*ean AUC = 0.52) in the left central region (*M*ean AUC = 0.54).

During the period for two groups: 6,010 ms to 6,207 ms for autistic adults and 6,002 ms to 6,270 ms for neurotypical adults, we observed the peak of one negative ERP at channels FC3, FC1, FCZ, FC2, FC4, C3, C1, CZ, C2, C4, CP3, CP1, CPZ, CP2, and CP4 (Figure 6). There was a significant interaction effect (*F* = 8.70, *p* = .004, 95%CI = [-0.39, −0.08]). Autistic adults exhibited a significantly larger negative deflection for the condition BG+FE+ (*M* = −1.23, *SD* = 0.55, β = - 0.70, *SE* = 0.06, *t* = −10.76, *p* < .001, 95%CI = [-0.83, −0.57]) compared to the condition BG+FE-(*M* = −0.53, *SD* = 0.40), the same pattern for neurotypical adults (BG+FE+: *M* = −0.95, *SD* = 0.44; BG+FE-: *M* = −0.61, *SD* = 0.41; β = - 0.34, *SE* = 0.06, *t* = −5.56, *p* < .001, 95%CI = [-0.46, −0.22]) but with smaller estimates. Moreover, autistic adults exhibited a larger negative deflection for the condition BG+FE+ than neurotypical adults (β = −0.28, *SE* = 0.9, *t* = −3.31, *p* = .001, 95%CI = [-0.45, −0.11]).

**Figure 6.**
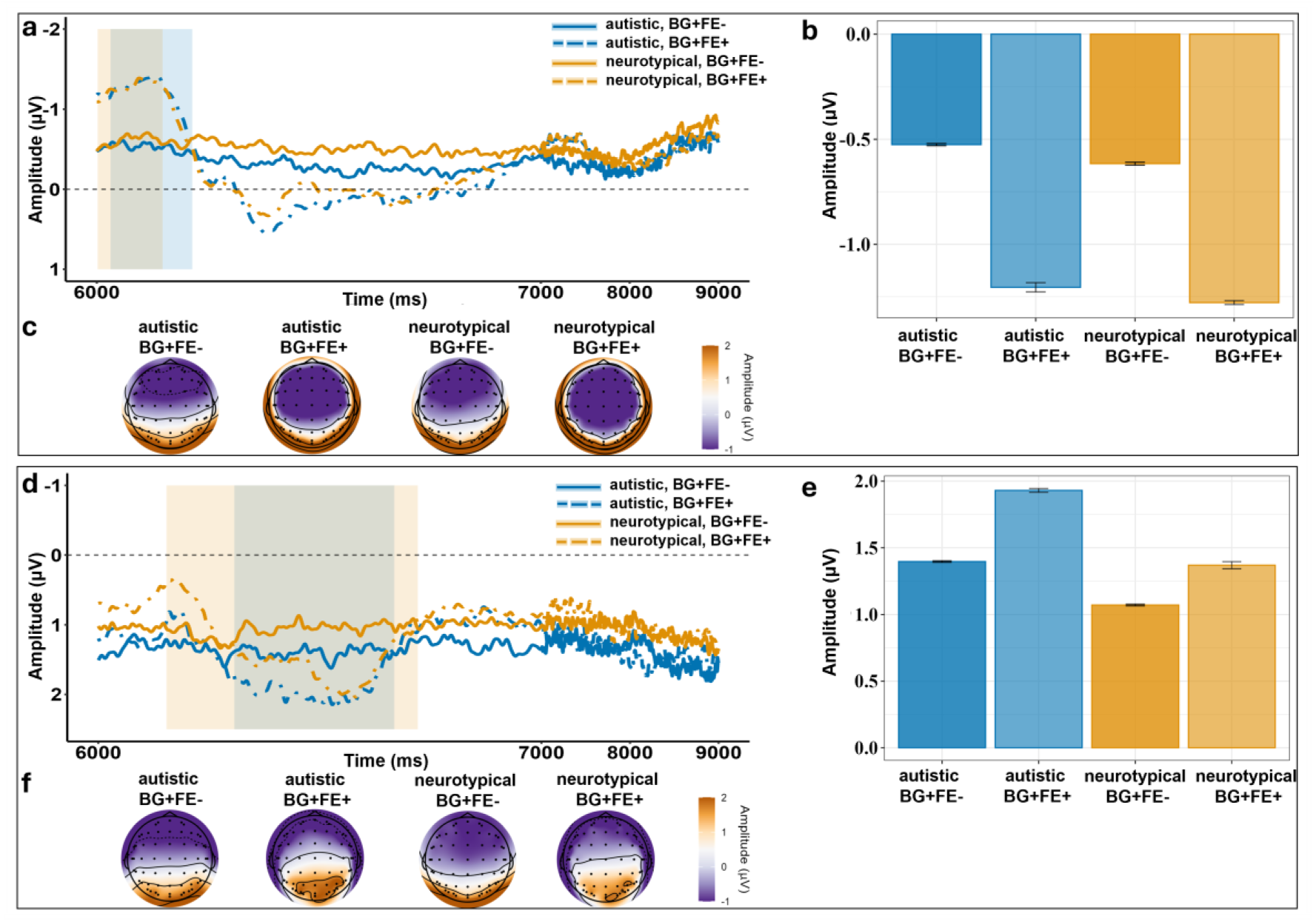
ERP Results Comparing BG+FE+ and BG+FE-During the Facial Expression Phase. *Note*. ERP waveforms display neural responses across two conditions and two groups over time, at the frontocentral, central, and centroparietal channels (Panel a), and at the parietal and parieto-occipital channels (Panel d). The topographic maps (Panel c) illustrate the amplitude distribution from 6,010 ms to 6,207 ms for autistic adults (highlighted by the light blue bar in Panel a) and from 6,002 ms to 6,270 ms for neurotypical adults (highlighted by the light yellow bar in Panel a). The topographic maps (Panel f) illustrate the amplitude distribution from 6,213 ms to 6,664 ms for autistic adults (highlighted by the light blue bar in Panel d) and from 6,283 ms to 6,775 ms for neurotypical adults (highlighted by the light yellow bar in Panel d). The bar charts (Panels b and e) compare averaged neural responses during corresponding time windows and at selected channels. The x-axis employs a piecewise linear transformation where the interval 7,000-9,000 ms is compressed for a better visualization.

During the period for two groups: 6,213 ms to 6,664 ms for autistic adults and 6,283 ms to 6,775 ms for neurotypical adults, we observed the peak of one positive ERP at channels P3, P1, PZ, P2, P4, PO3, POZ, and PO4 (Figure 6). Autistic adults (*M* = 1.62, *SD* = 0.98, β = 0.36, *SE* = 0.14, *t* = 2.66, *p* = .009, 95%CI = [0.09, 0.64]) exhibited significantly larger positive deflection than neurotypical adults (*M* = 1.26, *SD* = 0.79). The congruent condition (BG+FE+; *M* = 1.65, *SD* = 1.02, β = 0.44, *SE* = 0.09, *t* = 4.99, *p* < .001, 95%CI = [0.27, 0.61]) elicited significantly larger positive deflection than the congruent condition (BG+FE-; *M* = 1.21, *SD* = 0.70). No significant interaction effect was found (*p* > .05).

#### Comparing two congruent conditions (BG+FE+ vs. BG-FE-) between groups

Autistic adults exhibited significant above-channel classification in distinguishing pain (BG+FE+) versus no-pain events (BG-FE-), which sustained across time during two periods with channel-specific contributions (*p* clusters < .05, Figure 7a-c): (1) 6,031 ms-6,215 ms (*M*ean AUC = 0.55) in the temporal and frontocentral regions (*M*ean AUC = 0.54), and (2) 6,307 ms-6,668 ms (*M*ean AUC = 0.55) from the frontal to parietal regions (*M*ean AUC = 0.54).

**Figure 7.**
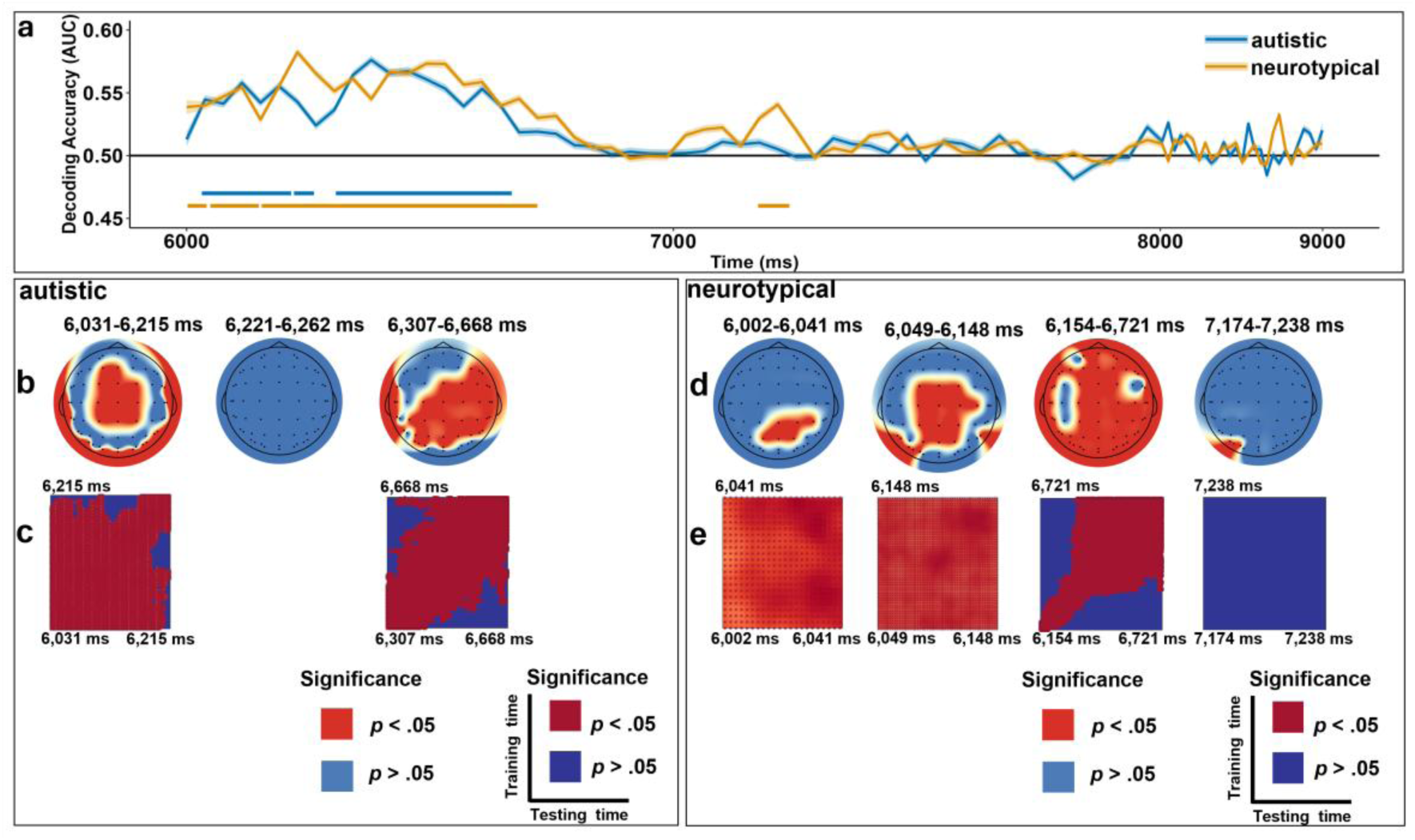
Temporal-based MVPA Results Comparing BG+FE+ and BG-FE-During the Facial Expression Phase. *Note*. Panel a: Classification accuracy in autistic (blue lines) and neurotypical (yellow lines) groups, where significant time clusters (*p* clusters < .05) are marked with horizontal bars below the decoding curves. Searchlight results for autistic (Panel b) and neurotypical (Panel d) groups reveal channels with (red dots, *p* clusters < .05) and without significant (blue dots, *p* clusters > .05) contributions during certain periods. Temporal generalization matrices for autistic (Panel c) and neurotypical (Panel e) groups mark periods with significant classification stability (*p* clusters < .05) in dark red, while unstable periods (*p* clusters > .05) in dark blue. The x-axis employs a piecewise linear transformation where the interval 8,000-9,000 ms is compressed for a better visualization.

Neurotypical adults showed significant classification sustaining across time during three periods with channel-specific contributions (*p* clusters < .05, Figure 7a, d, e): (1) 6,002 ms-6,041 ms (*M*ean AUC = 0.53) in the centroparietal region (*M*ean AUC = 0.54), (2) 6,049 ms-6,148 ms (*M*ean AUC = 0.55) from the frontal to parietal regions (*M*ean AUC = 0.54), and (3) 6,154 ms-6,721 ms (*M*ean AUC = 0.55) at almost the entire brain (*M*ean AUC = 0.54).

During the period for two groups: 6,031 ms to 6,215 ms for autistic adults and 6,002 ms to 6,148 ms for neurotypical adults, we observed the peak of one negative ERP at channels C3, C1, CZ, C2, C4, CP3, CP1, CPZ, CP2, CP4, P3, P1, PZ, P2, and P4 (Figure 8). The pain condition (BG+FE+; *M* = 0.05, *SD* = 0.56, β = −0.38, *SE* = 0.06, *t* = −6.15, *p* < .001, 95%CI = [-0.50, −0.26]) elicited significantly larger negative deflection than the no-pain condition (BG-FE-; *M* = 0.44, *SD* = 0.50). No significant group-relevant main or interaction effect was found (*p*s > .05).

**Figure 8.**
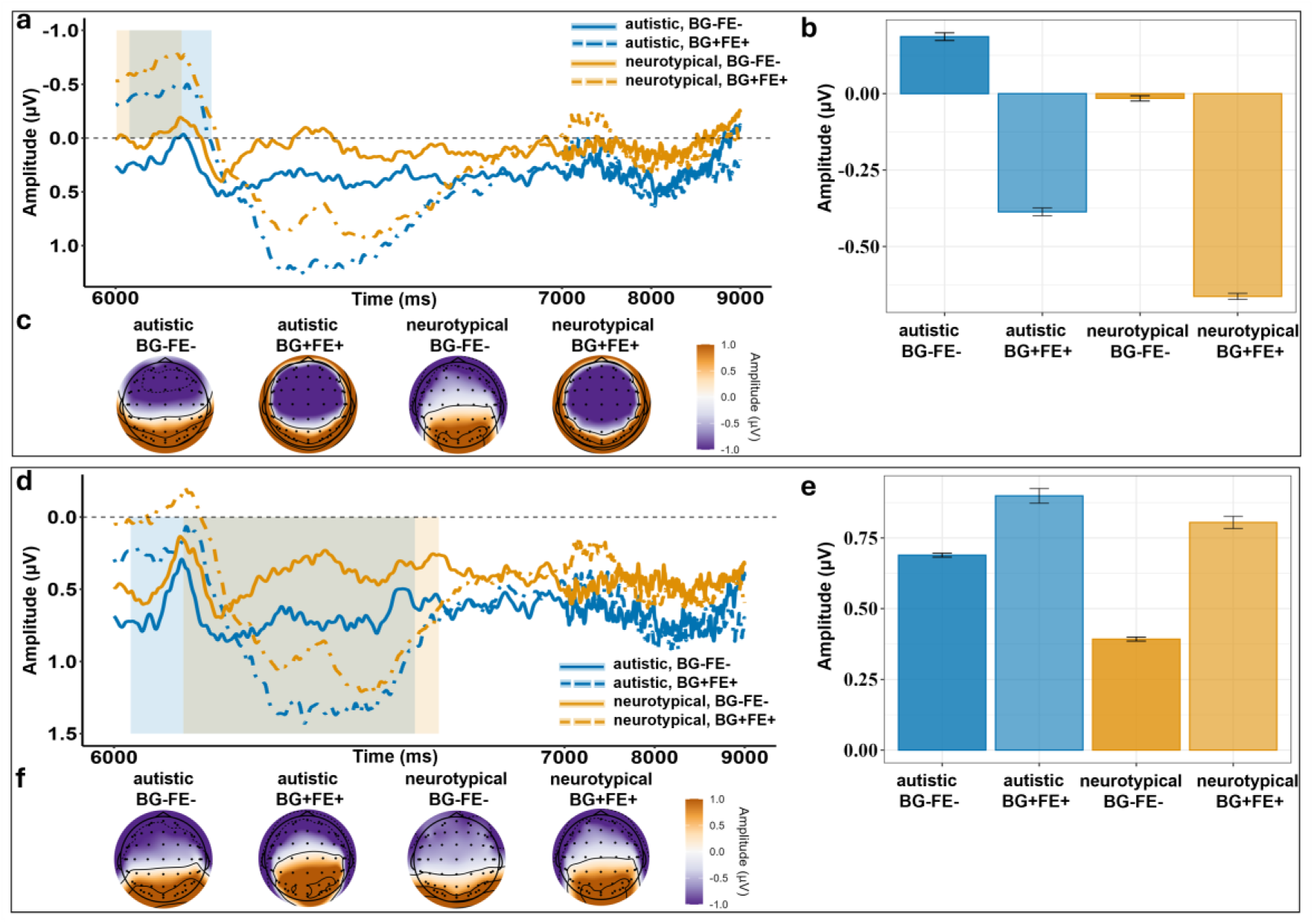
ERP Results Comparing BG+FE+ and BG-FE-During the Facial Expression Phase. *Note*. ERP waveforms display neural responses across two conditions and two groups over time, at the central, centroparietal, and parietal channels (Panel a), and the central, centroparietal, parietal, and parieto-occipital channels (Panel d). The topographic maps (Panel c) illustrate the amplitude distribution from 6,031 ms to 6,215 ms for autistic adults (highlighted by the light blue bar in Panel a) and from 6,002 ms to 6,148 ms for neurotypical adults (highlighted by the light yellow bar in Panel a). The topographic maps (Panel f) illustrate the amplitude distribution from 6,307 ms to 6,668 ms for autistic adults (highlighted by the light blue bar in Panel d) and from 6,154 ms to 6,721 ms for neurotypical adults (highlighted by the light yellow bar in Panel d). The bar charts (Panels b Panel e) compare averaged neural responses during corresponding time windows and at selected channels. The x-axis employs a piecewise linear transformation where the interval 7,000-9,000 ms is compressed for a better visualization.

Additionally, during the period for two groups: 6,307 ms to 6,668 ms for autistic adults and 6,154 ms to 6,721 ms for neurotypical adults, we observed the peak of one positive ERP at channels C3, C1, CZ, C2, C4, CP3, CP1, CPZ, CP2, CP4, P3, P1, PZ, P2, P4, PO5, PO3, POZ, PO4, and PO6 (Figure 8). There was a significant interaction effect (*F* = 5.76, *p* = .018, 95%CI = [0.05, 0.47]). Autistic adults exhibited a significantly larger positive deflection for the pain condition (*M* = 1.09, *SD* = 0.74, β = 0.71, *SE* = 0.08, *t* = 9.11, *p* < .001, 95%CI = [0.56, 0.87]) compared to the no-pain condition (*M* = 0.37, *SD* = 0.46), the same pattern for neurotypical adults (BG+FE+: *M* = 0.55, *SD* = 0.39; BG-FE-: *M* = 0.10, *SD* = 0.37; β = 0.46, *SE* = 0.07, *t* = 6.21, *p* < .001, 95%CI = [0.31, 0.60]) but with smaller estimates.

#### Comparing two incongruent conditions (BG+FE-vs. BG-FE+) between groups

Autistic adults exhibited significant above-channel classification in distinguishing two incongruent pain expressions (BG+FE-vs. BG-FE+), which sustained across time during three periods with channel-specific contributions (*p* clusters < .05, Figure 9a-c): (1) 6,064 ms-6,311 ms (*M*ean AUC = 0.55) in the frontotemporal and centroparietal regions (*M*ean AUC = 0.53), (2) 6,414 ms-6,455 ms (*M*ean AUC = 0.55) in the temporal and centroparietal regions (*M*ean AUC = 0.54), and (3) 6,461 ms-6,736 ms (*M*ean AUC = 0.54) in the centroparietal region (*M*ean AUC = 0.53).

**Figure 9.**
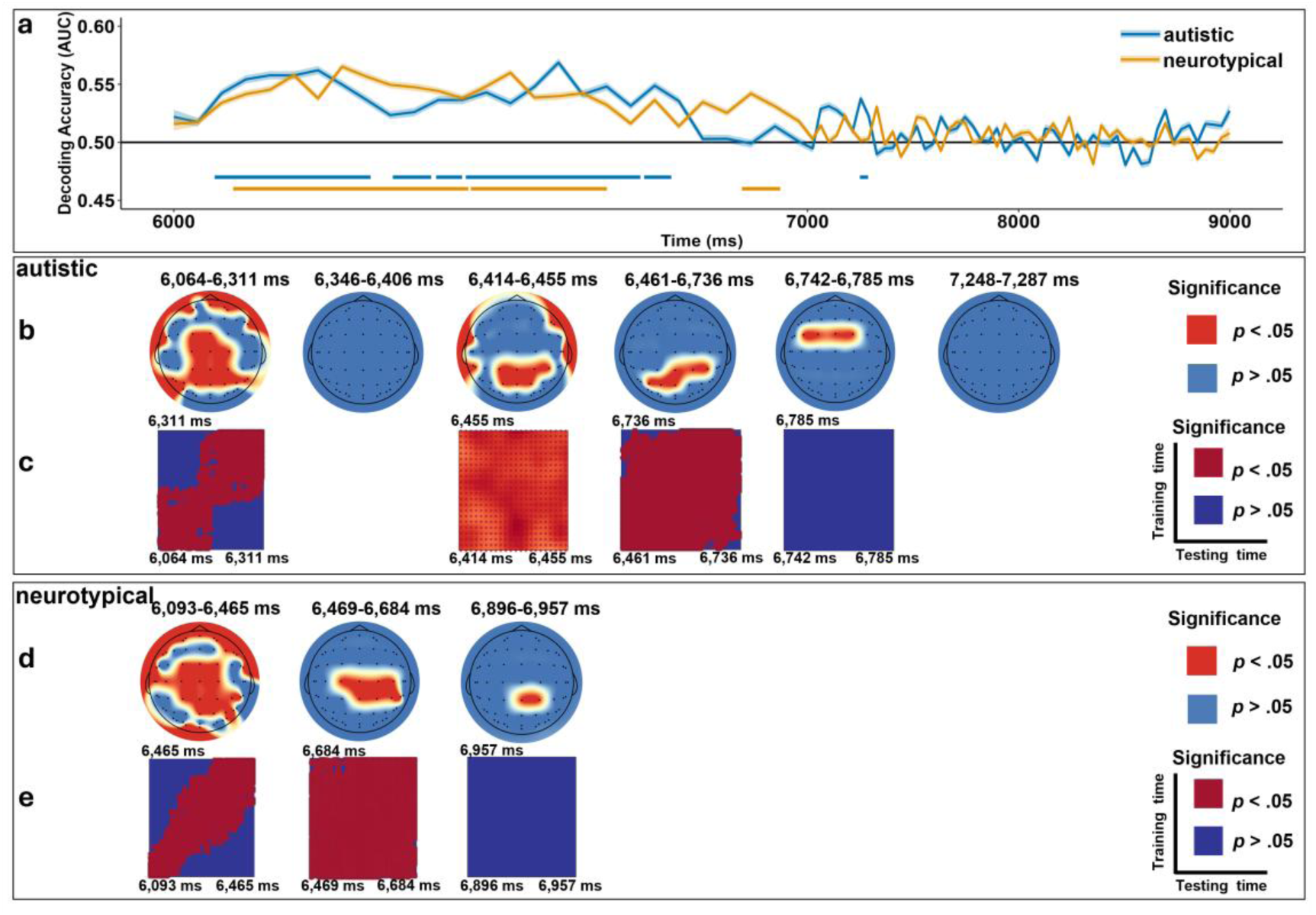
Temporal-based MVPA Results Comparing BG+FE- and BG-FE+ During the Facial Expression Phase. *Note*. Panel a: Classification accuracy in autistic (blue lines) and neurotypical (yellow lines) groups, where significant time clusters (*p* clusters < .05) are marked with horizontal bars below the decoding curves. Searchlight results for autistic (Panel b) and neurotypical (Panel d) groups reveal channels with (red dots, *p* clusters < .05) and without significant (blue dots, *p* clusters > .05) contributions during certain periods. Temporal generalization matrices for autistic (Panel c) and neurotypical (Panel e) groups mark periods with significant classification stability (*p* clusters < .05) in dark red, while unstable periods (*p* clusters > .05) in dark blue. The x-axis employs a piecewise linear transformation where the interval 7,000-9,000 ms is compressed for a better visualization.

Neurotypical adults exhibited significant classification, which sustained across time during two periods with channel-specific contributions (*p* clusters < .05, Figure 9a, d, e): (1) 6,093 ms-6,465 ms (*M*ean AUC = 0.55) from the anterior to occipital regions (*M*ean AUC = 0.53), and (2) 6469 ms-6,684 ms (*M*ean AUC = 0.54) in the centroparietal region (*M*ean AUC = 0.55).

We only observed the peak of one negative ERP for autistic adults between 6,064 ms and 6,311 ms at channels C3, C1, CZ, C2, C4, CP3, CP1, CPZ, CP2, CP4, P3, P1, PZ, P2, and P4 (Figure 10), with a significantly larger negative deflection for the condition BG-FE+ (*M* = −0.19, *SD* = 0.51, β = 0.46, *SE* = 0.07, *t* = 6.60, *p* < .001, 95%CI = [0.32, 0.60]) than BG+FE-(*M* = 0.27, *SD* = 0.43).

**Figure 10.**
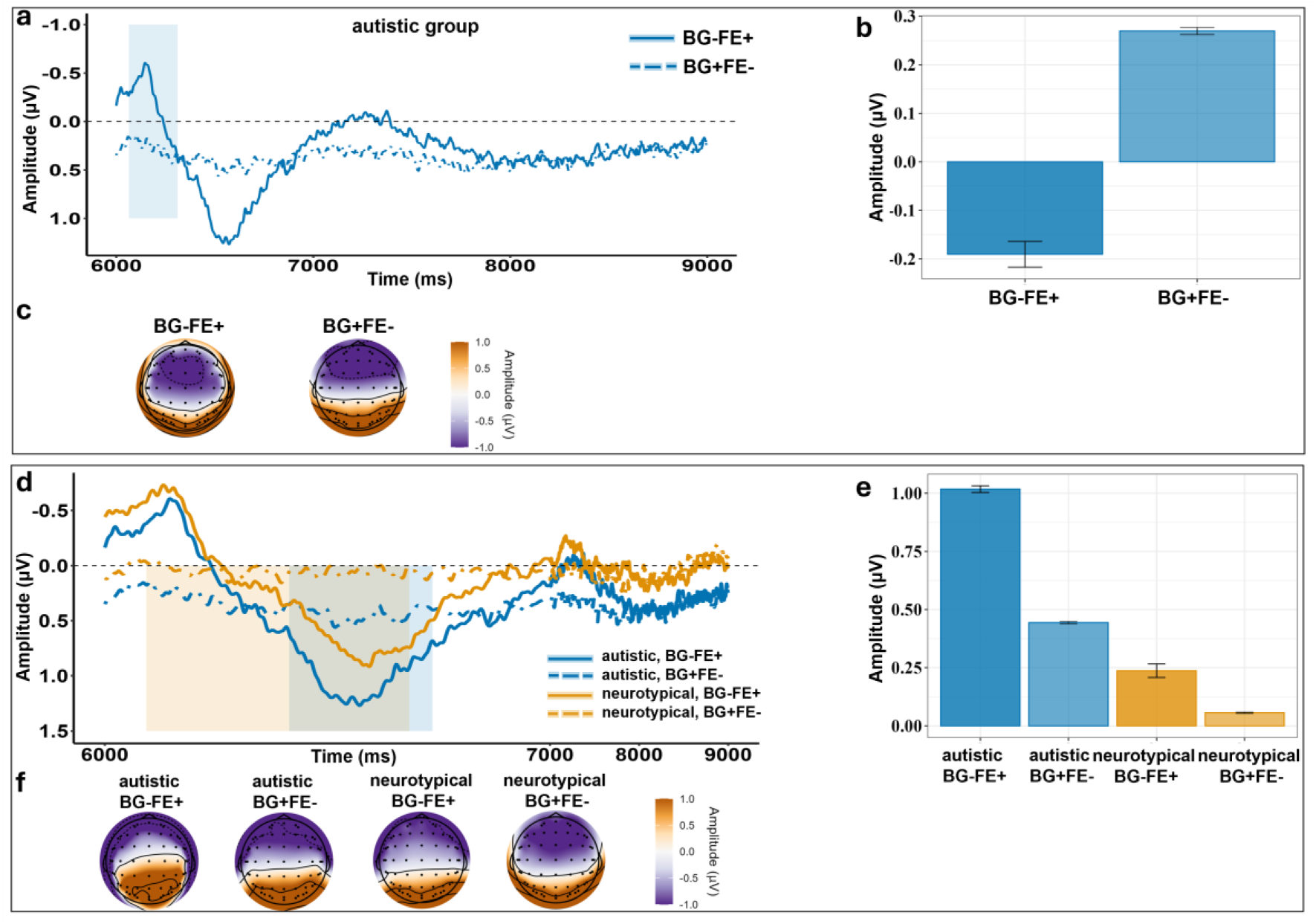
ERP Results Comparing BG+FE- and BG-FE+ During the Facial Expression Phase. *Note*. ERP waveforms display neural responses across two conditions over time at the central, centroparietal, and parietal channels for autistic adults (Panel a), and across two conditions and two groups at the central, centroparietal, and parietal channels (Panel d). The topographic maps (Panel c) illustrate the amplitude distribution from 6,093 ms to 6,465 ms for autistic adults (highlighted by the light blue bar in Panel a). The topographic maps (Panel f) illustrate the amplitude distribution from 6,414 ms to 6,736 ms for autistic adults (highlighted by the light blue bar in Panel d) and from 6,093 ms to 6,684 ms for neurotypical adults (highlighted by the light yellow bar in Panel d). The bar charts (Panel b) compare averaged neural responses during corresponding time windows and at selected channels for autistic adults (Panel b) and for both groups (Panel e). The x-axis employs a piecewise linear transformation where the interval 7,000-9,000 ms is compressed for a better visualization.

During the later period for two groups: 6,414 ms to 6,736 ms for autistic adults and 6,093 ms to 6,684 ms for neurotypical adults, we observed the peak of one positive ERP at channels C3, C1, CZ, C2, C4, CP3, CP1, CPZ, CP2, CP4, P3, P1, PZ, P2, and P4 (Figure 10). There was a significant interaction effect (*F* = 19.48, *p* < .001, 95%CI = [0.22, 0.57]). Autistic adults exhibited a significantly larger positive deflection for the condition BG-FE+(*M* = 1.02, *SD* = 0.57, β = 0.57, *SE* = 0.07, *t* = 8.75, *p* < .001, 95%CI = [0.44, 0.70]) than the condition BG+FE-(*M* = 0.44, *SD* = 0.46), the same pattern for neurotypical adults (BG+FE+: *M* = 0.26, *SD* = 0.24; BG-FE-: *M* = 0.09, *SD* = 0.13; β = 0.18, *SE* = 0.06, *t* = 2.88, *p* = .005, 95%CI = [0.06, 0.30]) but with smaller estimates.

### Multivariate and Univariate Analyses of Frequency-based EEG

#### Data

Results of frequency-based MVPA and univariate analysis were reported according to different phases and condition contrasts.

#### Body Gesture Phase

Both autistic and neurotypical adults exhibited significant above-chance classification when distinguishing body gestures in response to pain versus no-pain events, sustaining across time in the theta (3–8 Hz, Figure 11c, l), but not in the mu (8–13 Hz, Figure 11f, o) and beta (13–30 Hz, Figure 11i, r) bands.

**Figure 11.**
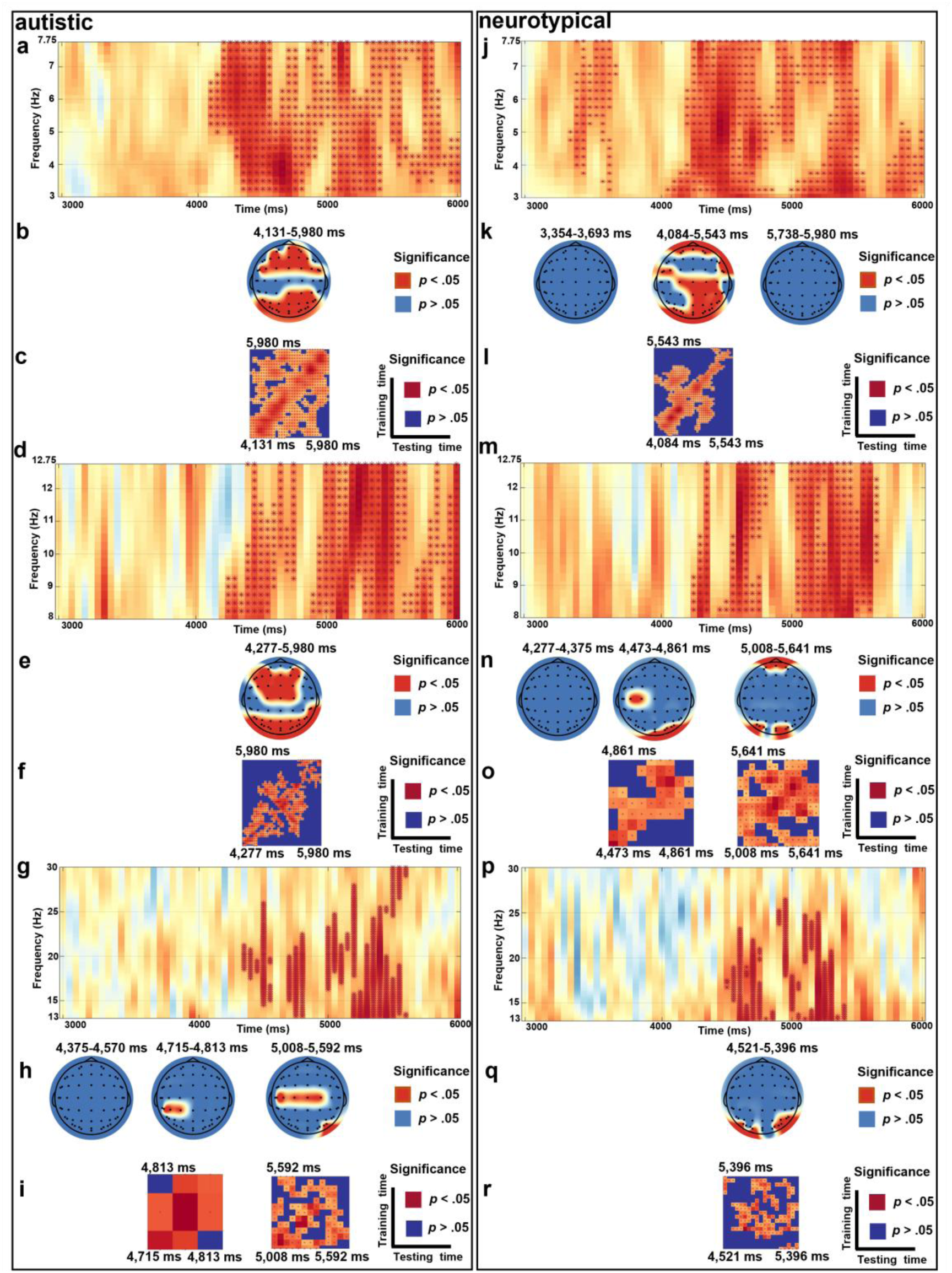
Frequency-based MVPA Comparing Pain versus No-pain During the Body Gesture Phase. *Note*. Left Panel: Autistic group; Right Panel: Neurotypical group. Classification accuracies for two groups are illustrated in three frequency bands, i.e., theta (Panels a and j), mu (Panels d and m), and beta (Panels g and p), where significant time-frequency clusters (*p* clusters < .05) are marked dark red asterisks. Searchlight results for two groups reveal channels with (red dots, *p* clusters < .05) and without significant (blue dots, *p* clusters > .05) contributions during certain periods in theta (Panels b and k), mu (Panels e and n), and beta (Panels h and q). Temporal generalization matrices for two groups mark periods with significant classification stability (*p* clusters < .05) in dark red, while unstable periods (*p* clusters > .05) in dark blue, in theta (Panels c and l), mu (Panels f and o), and beta (Panels i and r).

Specifically, autistic adults exhibited sustained significant classification between 4,131 ms and 5,980 ms (*p* clusters < .05, *M*ean AUC = 0.53, Figure 11a), particularly in the frontocentral and parieto-occipital regions (*p* clusters < .05, *M*ean AUC = 0.53, Figure 11b). Neurotypical adults exhibited sustained significance between 4,083 ms and 5,494 ms (*p* clusters < .05, *M*ean AUC = 0.53, Figure 11j), particularly in the centroparietal and parieto-occipital regions (*p* clusters < .05, *M*ean AUC = 0.52, Figure 11k).

During the period for two groups: 4,131 ms to 5,980 ms for autistic adults and 4,083 ms to 5,494 ms for neurotypical adults, we compared theta power at shared channels with significant contributions (Figure 12) but observed no significant group or condition relevant main or interaction effects (*p*s > .05).

**Figure 12.**
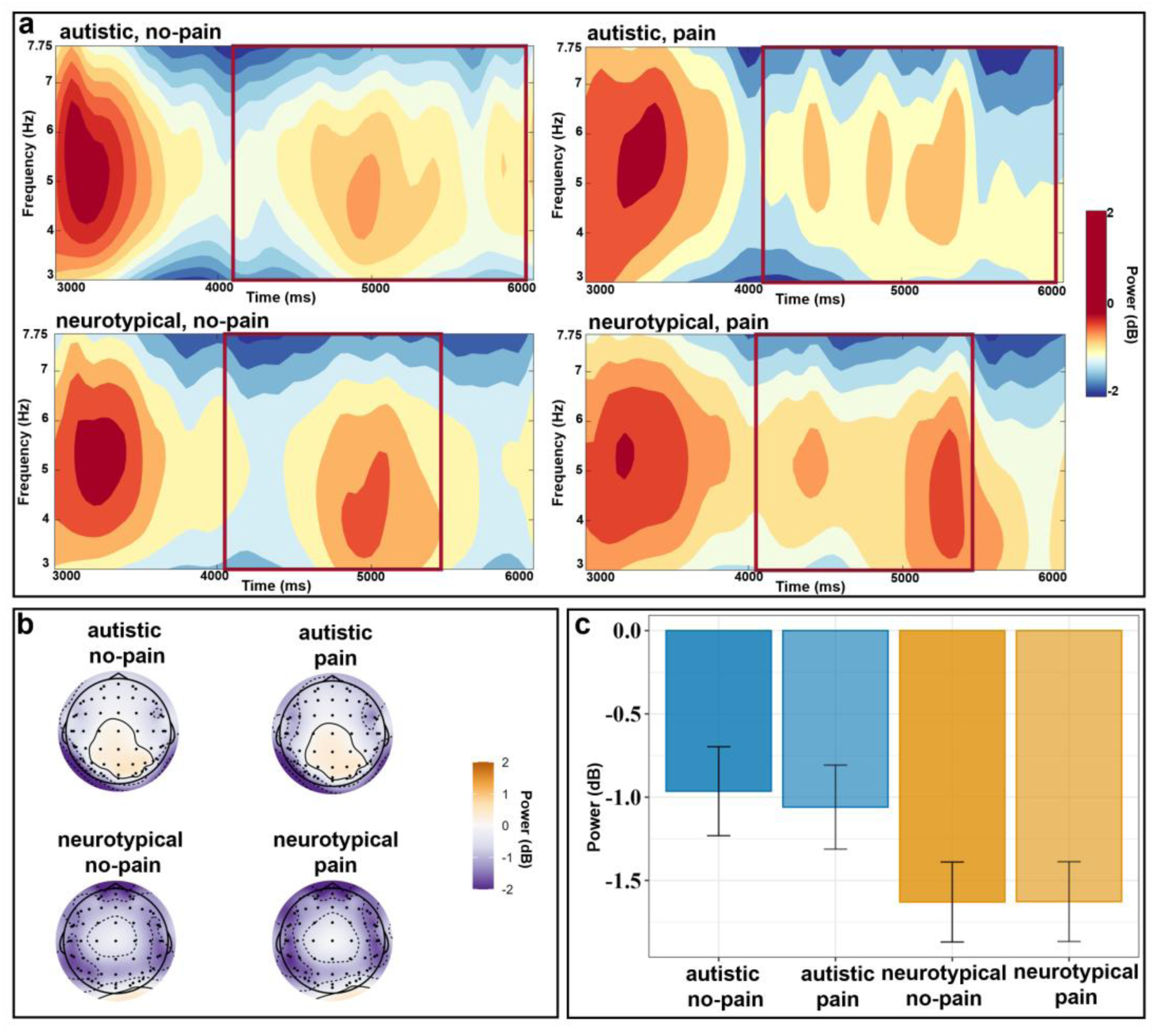
ERSP Comparing Pain versus No-pain During the Body Gesture Phase in the Theta Band. *Note*. Panel a: ERSP power distributions across two conditions for two groups over time at the frontal, frontocentral, centroparietal, parietal, parieto-occipital, and occipital channels. The topographic maps (Panel b) illustrate power distribution from 4,131 ms to 5,980 ms for autistic adults (highlighted by red rectangles in the first row of Panel a), and from 4,083 ms to 5,494 ms for neurotypical adults (highlighted by red rectangles in the second row of Panel a). The bar charts (Panel c) compare power between two conditions and groups during corresponding time windows and at selected channels.

### Facial Expression Phase

#### Comparing congruence between two pain expressions under no-pain events (BG-FE-vs. BG-FE+) between two groups

Both autistic and neurotypical adults exhibited significant above-chance classification, sustaining across time in the theta (3–8 Hz, Figure 13c, j), but not in the mu (8–13 Hz, Figure 13f, m) and beta (13–30 Hz, Figure 13g, n) bands.

**Figure 13.**
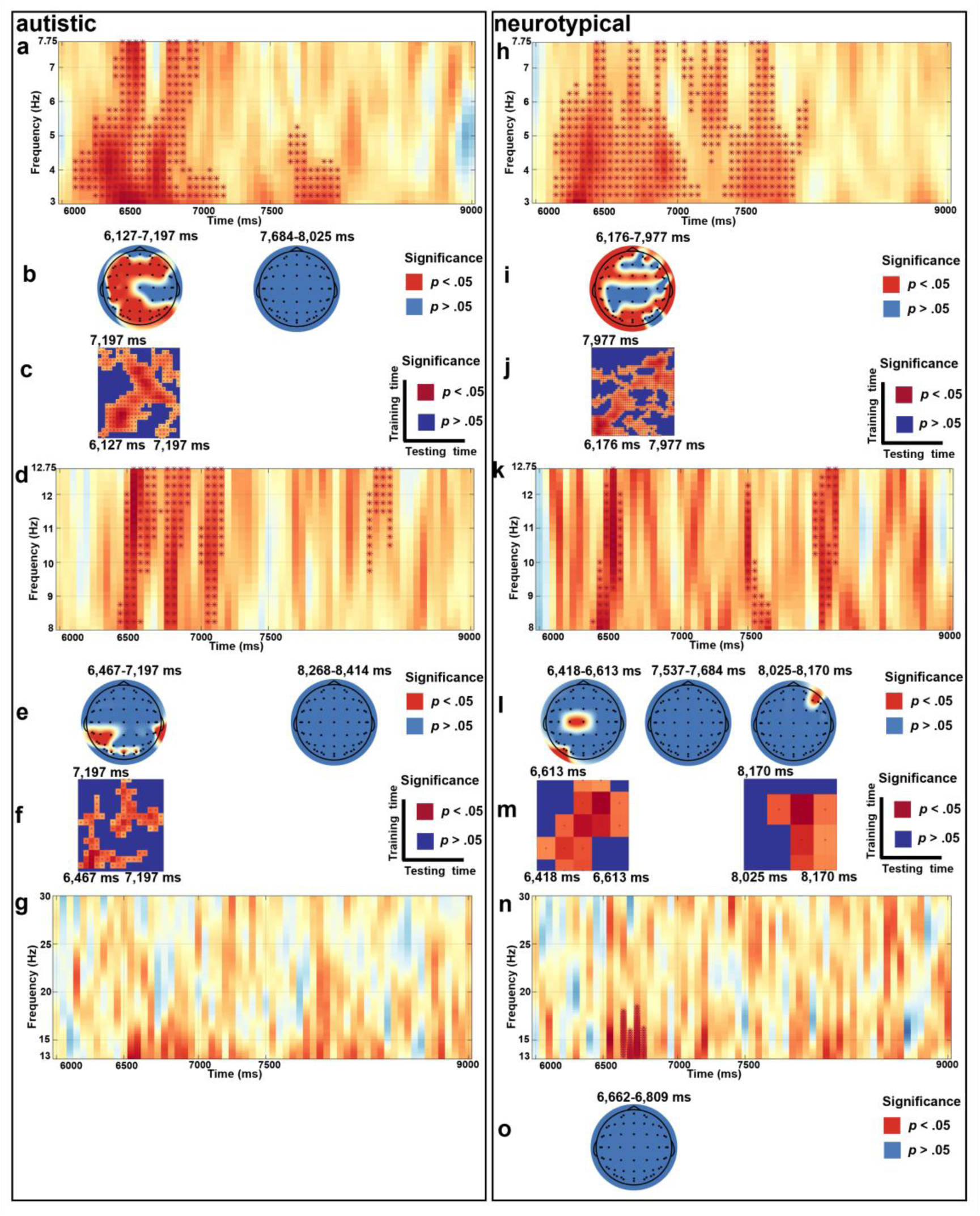
Frequency-based MVPA Comparing BG-FE- and BG-FE+ During the Facial Expression Phase. *Note*. Left Panel: Autistic group; Right Panel: Neurotypical group. Classification accuracies for two groups are illustrated in three frequency bands, i.e., theta (Panels a and h), mu (Panels d and k), and beta (Panels g and n), where significant time-frequency clusters (*p* clusters < .05) are marked dark red asterisks. Searchlight results for two groups reveal channels with (red dots, *p* clusters < .05) and without significant (blue dots, *p* clusters > .05) contributions during certain periods in theta (Panels b and i), mu (Panels e and l) for both groups, and beta (Panel o) only for neurotypical group. Temporal generalization matrices for two groups mark periods with significant classification stability (*p* clusters < .05) in dark red, while unstable periods (*p* clusters > .05) in dark blue, in theta (Panels c and j) and mu (Panels f and m).

Specifically, autistic adults exhibited sustained significant classification between 6,176 ms and 6,857 ms (*p* clusters < .05, *M*ean AUC = 0.54, Figure 13a), particularly in the frontocentral and parieto-occipital regions (*p* clusters < .05, *M*ean AUC = 0.54, Figure 13b). Neurotypical adults exhibited sustained significance between 6,176 ms and 6,955 ms (*p* clusters < .05, *M*ean AUC = 0.54, Figure 13h), particularly in the frontocentral and parieto-occipital regions (*p* clusters < .05, *M*ean AUC = 0.54, Figure 13i).

During the period for two groups: 6,176 ms to 6,857 ms for autistic adults and 6,176 ms to 6,955 ms for neurotypical adults, we compared theta power at common channels with significant contributions: AF3, AF4, F7, F5, F3, FT7, FC5, FC3, FC1, FCZ, FC2, FC4, C5, CP5, P1, PZ, P2, P4, P8, PO7, PO5, PO3, POZ, O1, OZ, and O2 (Figure 14). Autistic adults (*M* = −1.00, *SD* = 1.85, β = 0.70, *SE* = 0.35, *t* = 2.01, *p* = .047, 95%CI = [0.01, 1.39]) induced significantly larger theta power than neurotypical adults (*M* = −1.70, *SD* = 1.82). The incongruent condition (BG-FE+; *M* = −1.14, *SD* = 1.82, β = 0.45, *SE* = 0.05, *t* = 9.85, *p* < .001, 95%CI = [0.36, 0.55]) elicited significantly larger theta power than the congruent condition (BG-FE-; *M* = - 1.60, *SD* = 1.88). No significant interaction effect was found (*p* > .05).

**Figure 14.**
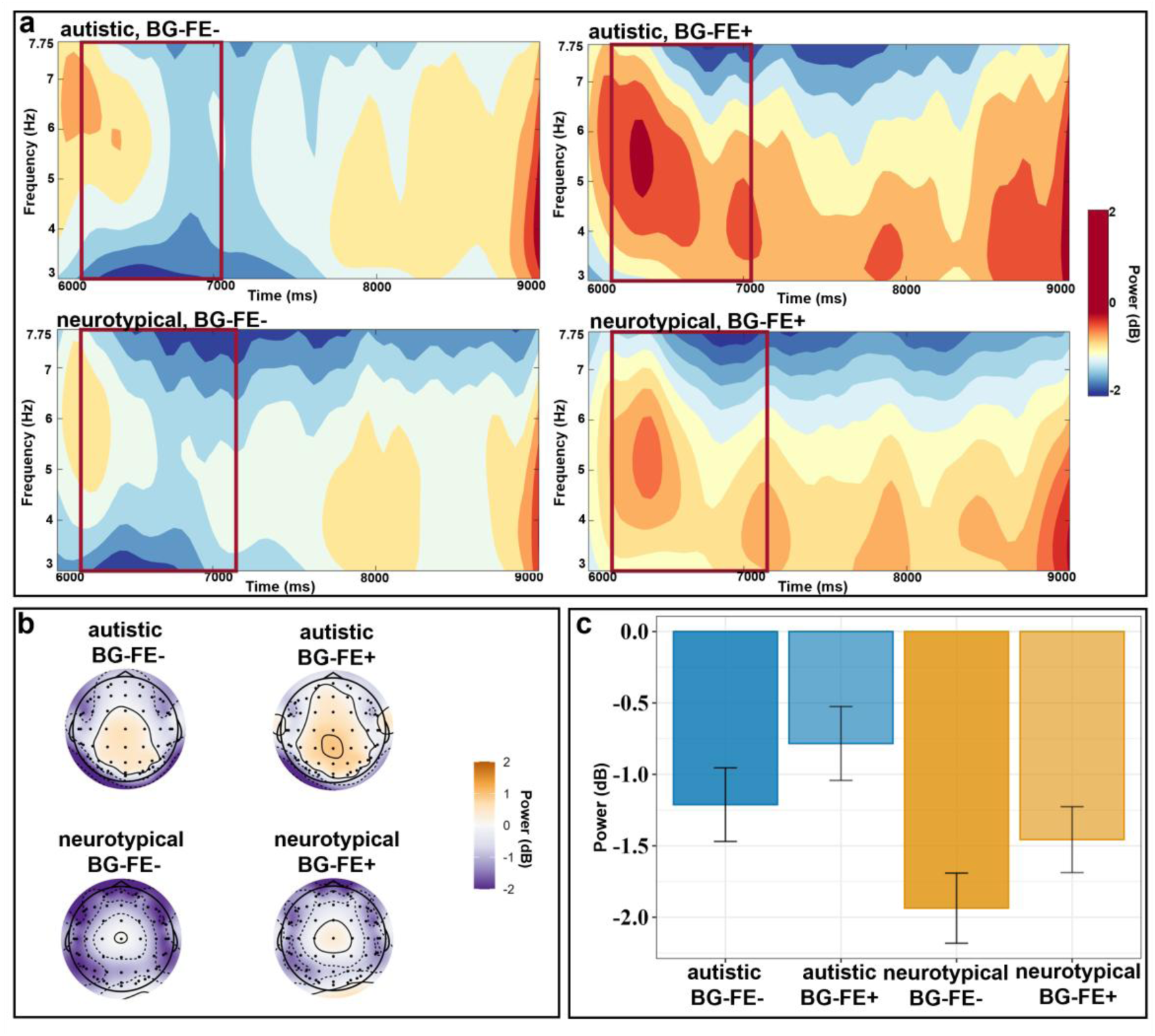
ERSP Comparing BG-FE- and BG-FE+ During the Facial Expression Phase in the Theta Band. *Note*. Panel a: ERSP power distributions across two conditions for two groups over time from the frontal to occipital channels. The topographic maps (Panel b) illustrate power distribution from 6,176 ms to 6,857 ms (highlighted by red rectangles in the first row of Panel a) for autistic group, and from 6,176 ms to 6,955 ms (highlighted by red rectangles in the second row of Panel a) for neurotypical group. The bar charts (Panel c) compare power between two conditions and two groups during corresponding time windows and at selected channels.

#### Comparing congruence between two pain expressions under no-pain events (BG+FE+ vs. BG+FE-) between two groups

Both autistic and neurotypical adults exhibited significant above-chance classification accuracy, sustaining across time in the theta (3–8 Hz, Figure 15c, l) and mu (8–13 Hz, Figure 15f, o) bands, but not in the beta (13–30 Hz, Figure 15i, r) band.

**Figure 15.**
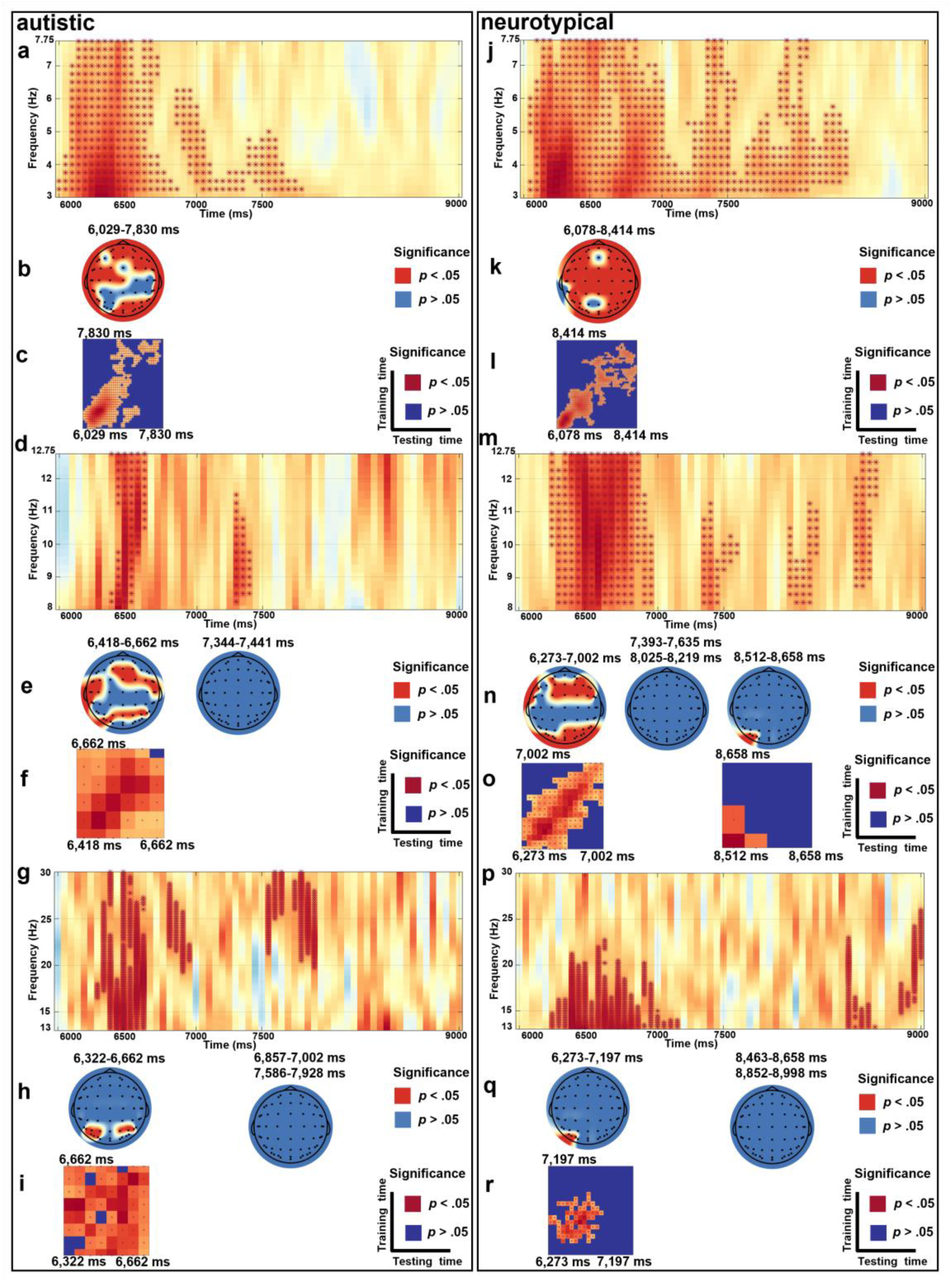
Frequency-based MVPA Comparing BG+FE+ and BG+FE-During the Facial Expression Phase. *Note*. Left Panel: Autistic group; Right Panel: Neurotypical group. Classification accuracies for two groups are illustrated in three frequency bands: theta (Panels a and j), mu (Panels d and m), and beta (Panels g and p), where significant time-frequency clusters (*p* clusters < .05) are marked dark red asterisks. Searchlight results for two groups reveal channels with (red dots, *p* clusters < .05) and without significant (blue dots, *p* clusters > .05) contributions during certain periods in theta (Panels b and k), mu (Panels e and n), and beta (Panels h and q). Temporal generalization matrices for two groups mark periods with significant classification stability (*p* clusters < .05) in dark red, while unstable periods (*p* clusters > .05) in dark blue, in theta (Panels c and l), mu (Panels f and o), and beta (Panels i and r).

In the theta band (3–8 Hz), autistic adults exhibited sustained significant classification between 6,127 ms and 7,100 ms (*p* clusters < .05, *M*ean AUC = 0.55, Figure 15a), particularly in the frontocentral and parieto-occipital regions (*p* clusters < .05, *M*ean AUC = 0.54, Figure 15b). Neurotypical adults exhibited sustained significant classification between 6,078 ms and 7,051 ms (*p* clusters < .05, *M*ean AUC = 0.55, Figure 15j), at almost the entire brain (*p* clusters < .05, *M*ean AUC = 0.54, Figure 15k).

During the period for two groups: 6,127 ms to 7,100 ms for autistic adults and 6,078 ms to 7,051 ms for neurotypical adults, we compared theta power at common channels with significant contributions: FP1, FPZ, FP2, AF3, AF4, F5, F1, F2, F4, F6, F8, FT7, FC5, FC3, FC1, FC2, FC4, FC6, FT8, T7, C5, C3, C1, CZ, T8, CP5, TP8, P7, P5, P2, P4, P6, P8, POZ, PO4, PO6, PO8, O1, OZ, and O2 (Figure 16). The congruent condition (BG+FE+; *M* = −1.40, *SD* = 2.03, β = 0.46, *SE* = 0.05, *t* = 8.47, *p* < .001, 95%CI = [0.35, 0.57]) elicited significantly larger theta power than the incongruent condition (BG+FE-; *M* = −1.86, *SD* = 2.06). No significant group related main effect or interaction effect was found (*p* > .05).

**Figure 16.**
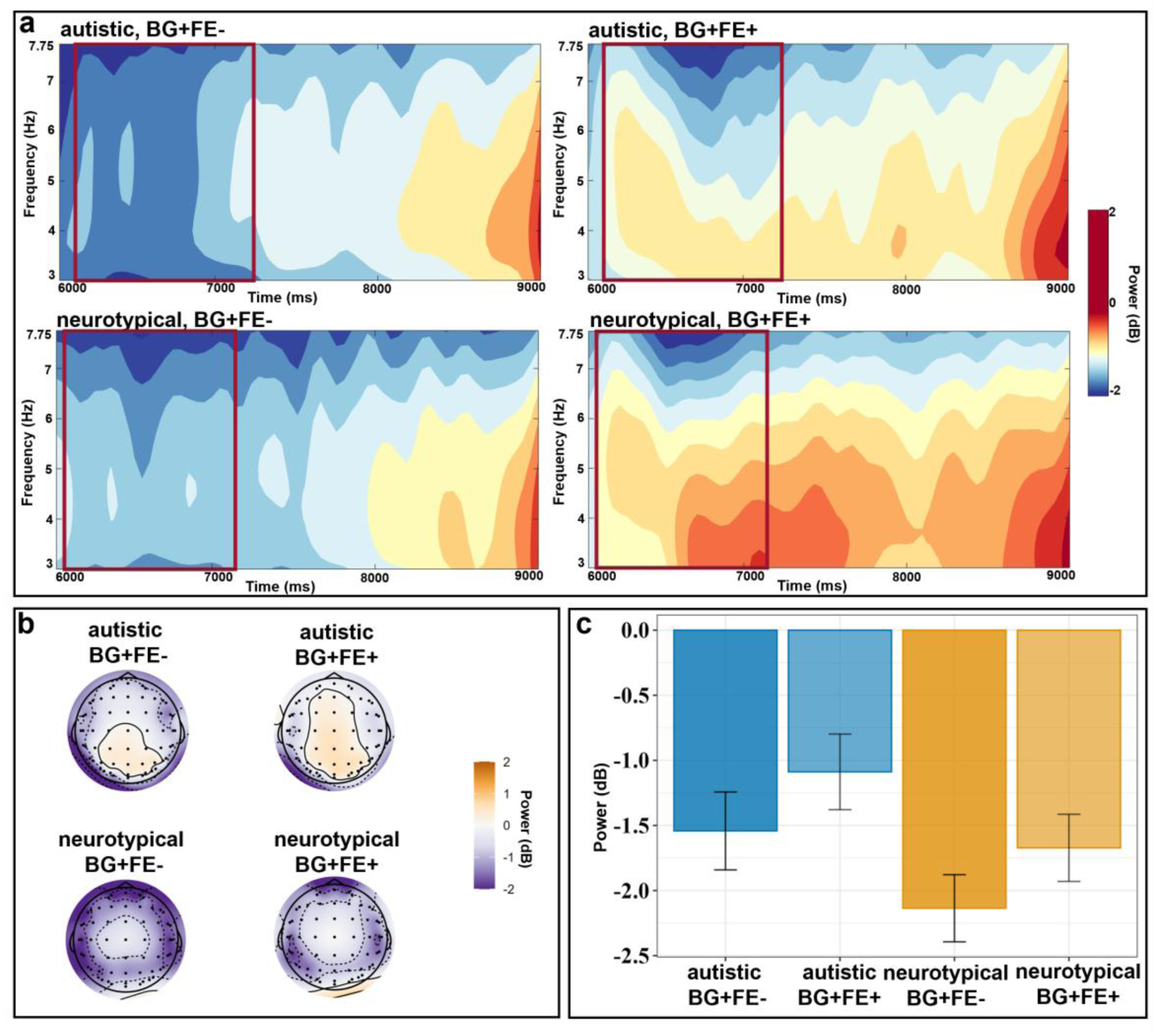
ERSP Comparing BG+FE+ and BG+FE-During the Facial Expression Phase in the Theta Band. *Note*. Panel a: ERSP power distributions across two conditions for two groups over time from the frontal to occipital channels. The topographic maps (Panel b) illustrate power distribution from 6,127 ms to 7,100 ms for autistic adults (highlighted by red rectangles in the first row of Panel a), and from 6,078 ms to 7,051 ms for neurotypical adults (highlighted by red rectangles in the second row of Panel a). The bar charts (Panel c) compare power between two conditions and two groups during corresponding time windows and at selected channels.

In the mu band (8–13 Hz), autistic adults exhibited sustained significant classification between 6,418 ms and 6,613 ms (*p* clusters < .05, *M*ean AUC = 0.54, Figure 15d), particularly in the frontocentral and parietal regions (*p* clusters < .05, *M*ean AUC = 0.53, Figure 15e). Neurotypical adults exhibited sustained significant classification between 6,273 ms and 7,002 ms (*p* clusters < .05, *M*ean AUC = 0.54, Figure 15m), particularly in the frontocentral and parieto-occipital regions (*p* clusters < .05, *M*ean AUC = 0.54, Figure 15n).

During the period for two groups: 6,418 ms to 6,613 ms for autistic adults and 6,273 ms to 7,002 ms for neurotypical adults, we compared theta power at common channels with significant contributions: F1, FZ, F2, FT7, FCZ, FC2, FC4, FC6, PZ, P2, P4, P6, P8, PO7, PO5, and PO3 (Figure 17). Neurotypical adults (*M* = −3.19, *SD* = 2.36, β = 1.03, *SE* = 0.43, *t* = 2.40, *p* = .018, 95%CI = [0.18, 1.87]) induced significantly larger suppression than autistic adults (*M* = - 2.17, *SD* = 2.12). The congruent condition (BG+FE+; *M* = −2.89, *SD* = 2.21, β = - 0.35, *SE* = 0.06, *t* = −6.00, *p* < .001, 95%CI = [-0.47, −0.24]) elicited significantly larger mu suppression than the congruent condition (BG+FE-; *M* = −2.53, *SD* = 2.38). No significant interaction effect was found (*p* > .05).

**Figure 17.**
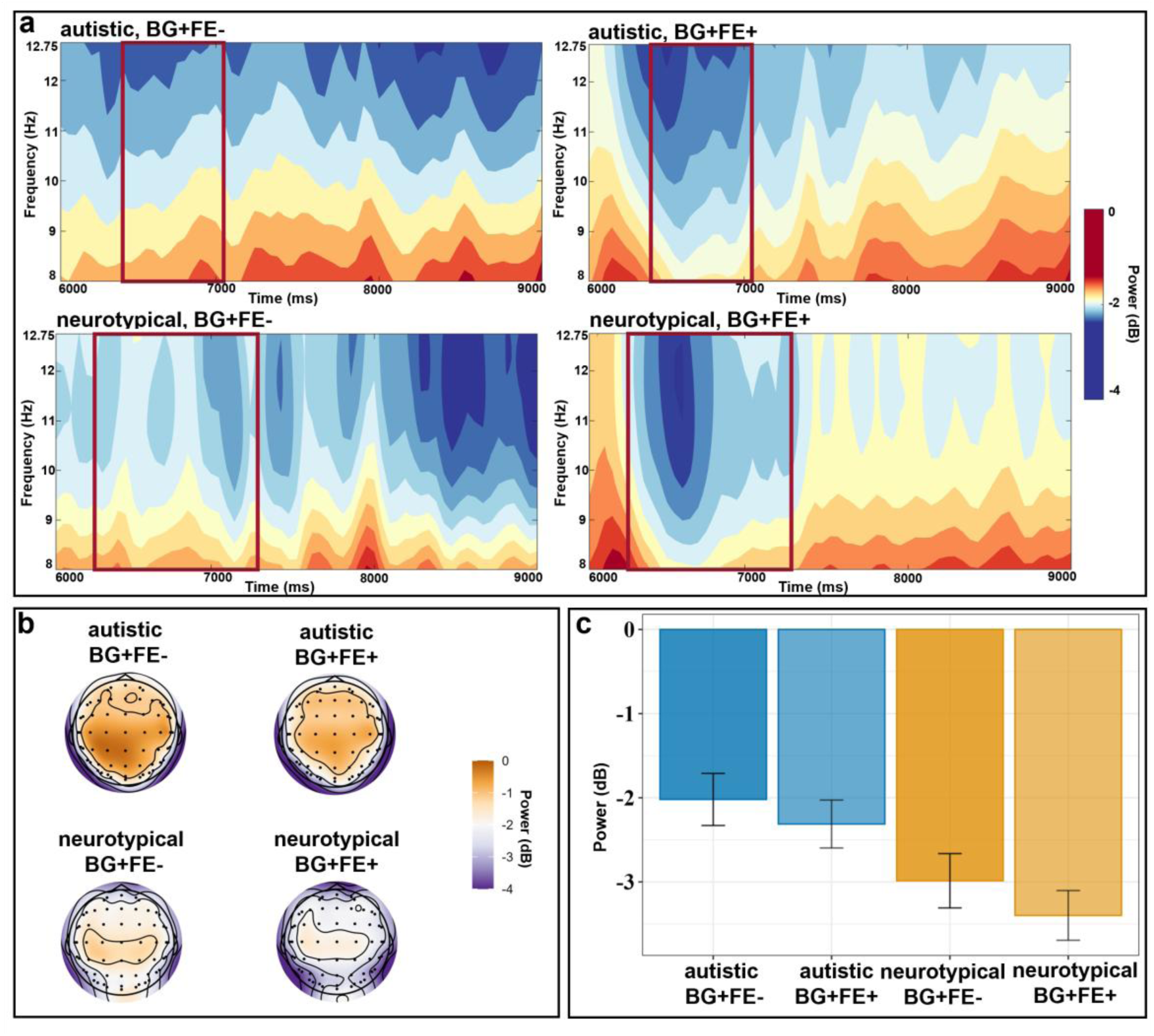
ERSP Comparing BG+FE+ and BG+FE-During the Facial Expression Phase in the Mu Band. *Note*. Panel a: ERSP power distributions across two conditions for two groups over time at the frontal, frontocentral, parietal, and parieto-occipital channels. The topographic maps (Panel b) illustrate power distribution from 6,418 ms to 6,613 ms for autistic adults (highlighted by red rectangles in the first row of Panel a), and from 6,273 ms to 7,002 ms for neurotypical adults (highlighted by red rectangles in the second row of Panel a). The bar charts (Panel c) compare power between two conditions and two groups during corresponding time windows and at selected channels.

#### Comparing two congruent conditions (BG+FE+ vs. BG-FE-)

Both autistic and neurotypical adults exhibited significant above-chance classification, sustaining across time in the theta (3–8 Hz, Figure 18c, l) band, but not in the mu (8–13 Hz, Figure 18f, o) and beta (13–30 Hz, Figure 18i, r) bands.

**Figure 18.**
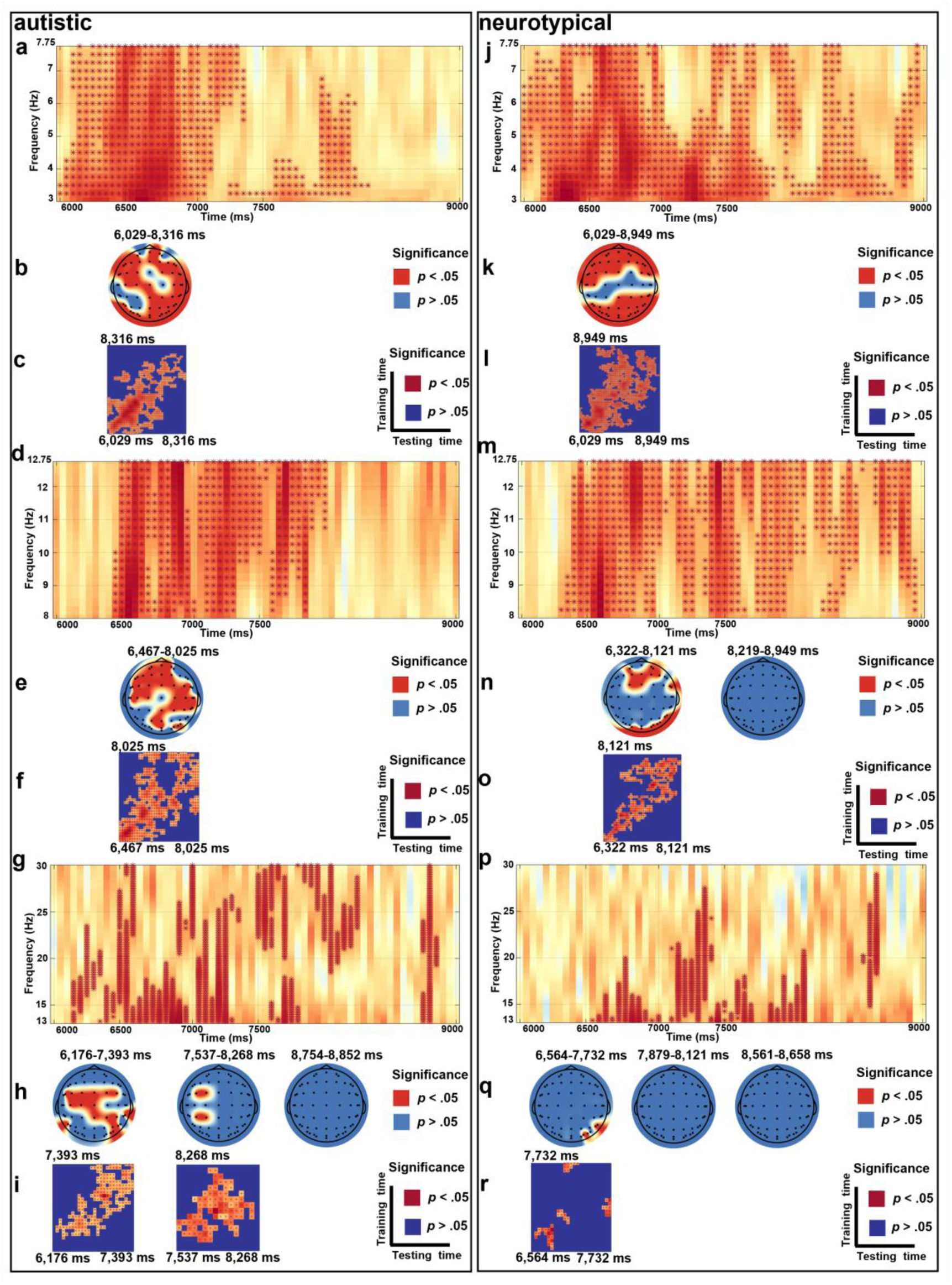
Frequency-based MVPA Comparing BG-FE- and BG+FE+ During the Facial Expression Phase. *Note*. Left Panel: Autistic group; Right Panel: Neurotypical group. Classification accuracies for two groups are illustrated in three frequency bands: theta (Panels a and j), mu (Panels d and m), and beta (Panels g and p), where significant time-frequency clusters (*p* clusters < .05) are marked dark red asterisks. Searchlight results for two groups reveal channels with (red dots, *p* clusters < .05) and without significant (blue dots, *p* clusters > .05) contributions during certain periods in theta (Panels b and k), mu (Panels e and n), and beta (Panels h and q). Temporal generalization matrices for two groups mark periods with significant classification stability (*p* clusters < .05) in dark red, while unstable periods (*p* clusters > .05) in dark blue, in theta (Panels c and l), mu (Panels f and o), and beta (Panels i and r).

In the theta band (3–8 Hz), autistic adults exhibited sustained significance between 6,078 ms and 7,002 ms (*p* clusters < .05, *M*ean AUC = 0.55, Figure 18a), particularly at most the entire brain (*p* clusters < .05, *M*ean AUC = 0.54, Figure 18b). Neurotypical adults exhibited sustained significance between 6,176 ms and 7,100 ms (*p* clusters < .05, *M*ean AUC = 0.54, Figure 18j) in the frontocentral and parieto-occipital regions (*p* clusters < .05, *M*ean AUC = 0.54, Figure 18k).

During the period for two groups: 6,078 ms to 7,002 ms for autistic adults and 6,176 ms to 7,100 ms for neurotypical adults, we compared theta power at common channels with significant contributions: FPZ, FP2, AF3, F7, F5, F3, F1, FZ, F2, F4, F6, F8, FT7, FC5, FC3, FC1, FC4, FC6, FT8, C3, CPZ, CP2, CP4, CP6, TP8, PZ, P2, P4, P6, P8, PO7, P5, PO3, POZ, PO4, PO6, PO8, O1, OZ, and O2 (Figure 19). The pain condition (BG+FE+; *M* = −1.29, *SD* = 2.01, β = 0.33, *SE* = 0.06, *t* = 6.00, *p* < .001, 95%CI = [0.22, 0.44]) elicited significantly larger theta power than the no-pain condition (BG-FE-; *M* = −1.63, *SD* = 1.91). No significant group relevant main effect or interaction effect was found (*p* > .05).

**Figure 19.**
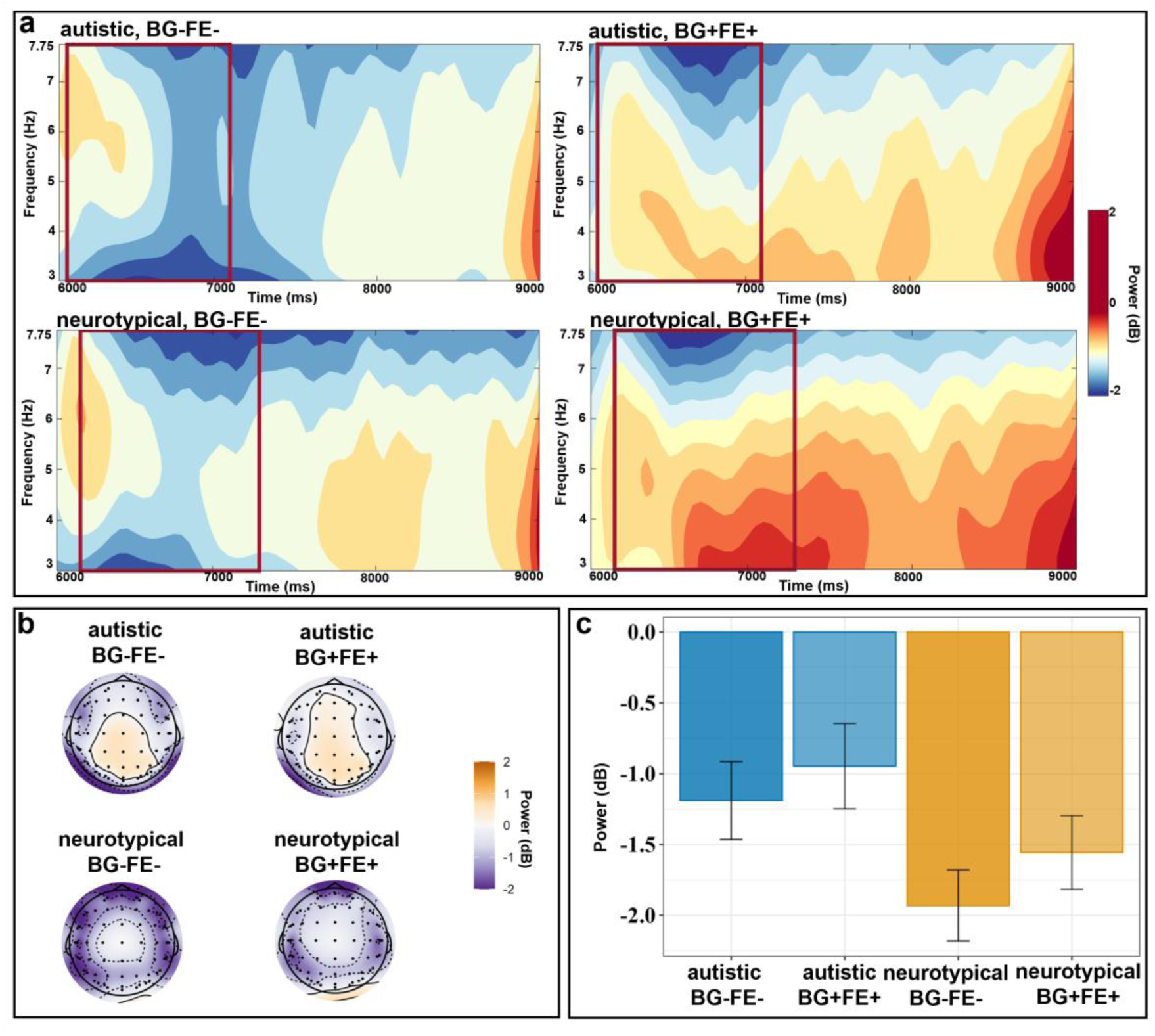
ERSP Comparing BG-FE- and BG+FE+ During the Facial Expression Phase in the Theta Band. *Note*. Panel a: ERSP power distributions across two conditions for two groups over time from the frontal to occipital channels. The topographic maps (Panel b) illustrate power distribution from 6,078 ms to 7,002 ms for autistic adults (highlighted by red rectangles in the first row of Panel a), and from 6,176 ms to 7,100 ms for neurotypical adults (highlighted by red rectangles in the second row of Panel a). The bar charts (Panel c) compare power between two conditions and two groups during corresponding time windows and at selected channels.

#### Comparing two incongruent conditions (BG+FE-vs. BG-FE+)

Both autistic and neurotypical adults exhibited significant above-chance classification accuracy when distinguishing body gestures in response to pain versus no-pain events, sustaining across time in the theta (3–8 Hz, Figure 20c, l) and mu (8–13 Hz, Figure 20f, o) bands, but not in the beta (13–30 Hz, Figure 20i, r) band.

**Figure 20.**
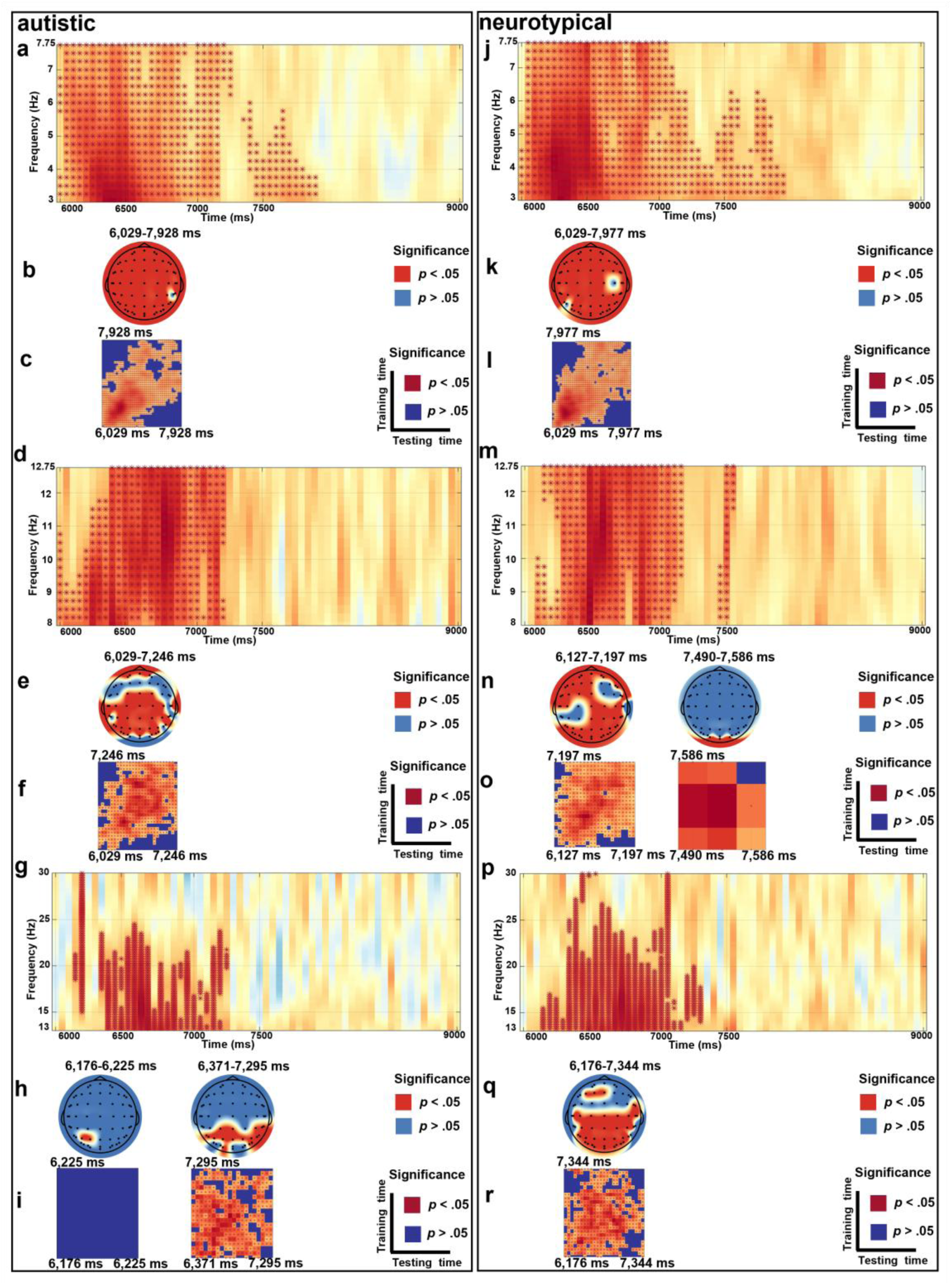
Frequency-based MVPA Comparing BG-FE+ and BG+FE-During the Facial Expression Phase. *Note*. Left Panel: Autistic group; Right Panel: Neurotypical group. Classification accuracies for two groups are illustrated in three frequency bands: theta (Panels a and j), mu (Panels d and m), and beta (Panels g and p), where significant time-frequency clusters (*p* clusters < .05) are marked dark red asterisks. Searchlight results for two groups reveal channels with (red dots, *p* clusters < .05) and without significant (blue dots, *p* clusters > .05) contributions during certain periods in theta (Panels b and k), mu (Panels e and n), and beta (Panels h and q). Temporal generalization matrices for two groups mark periods with significant classification stability (*p* clusters < .05) in dark red, while unstable periods (*p* clusters > .05) in dark blue, in theta (Panels c and l), mu (Panels f and o), and beta (Panels i and r).

In the theta band (3–8 Hz), autistic adults exhibited sustained significant classification between 6,078 ms and 7,197 ms (*p* clusters < .05, *M*ean AUC = 0.56, Figure 20a) in the most brain region (*p* clusters < .05, *M*ean AUC = 0.55, Figure 20b). Neurotypical adults exhibited sustained significance between 6,029 ms and 7,148 ms (*p* clusters < .05, *M*ean AUC = 0.55, Figure 20j) in the most brain region (*p* clusters < .05, *M*ean AUC = 0.54, Figure 20k).

During the period for two groups: 6,078 ms to 7,197 ms for autistic adults and 6,029 ms to 7,148 ms for neurotypical adults, we compared theta power at all 60 channels, except for C4, CP6, P5, with significant contributions (Figure 21). The incongruent condition under no-pain events (BG-FE+; *M* = −1.21, *SD* = 1.82, β = 0.56, *SE* = 0.06, *t* = 10.16, *p* < .001, 95%CI = [0.45, 0.67]) elicited significantly larger theta power than the incongruent condition under pain events (BG+FE-; *M* = −1.77, *SD* = 2.00). No significant group relevant main effect or interaction effect was found (*p* > .05).

**Figure 21.**
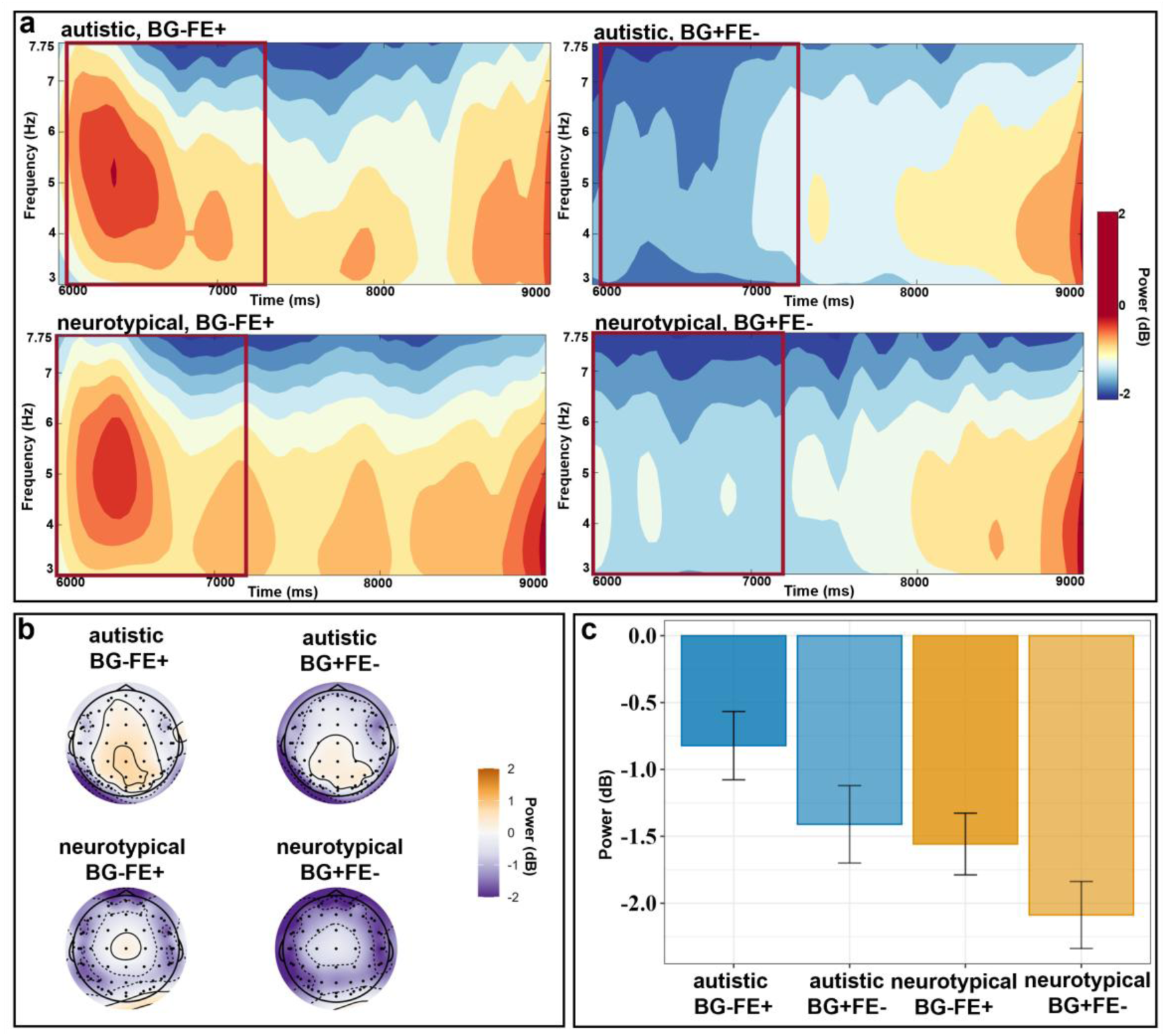
ERSP Comparing BG-FE+ and BG+FE-During the Facial Expression Phase in the Theta Band. *Note*. Panel a: ERSP power distributions across two conditions for two groups over time at all 60 channels, except for C4, CP6, P5. The topographic maps (Panel b) illustrate power distribution from 6,078 ms to 7,197 ms for autistic adults (highlighted by red rectangles in the first row of Panel a), and from 6,029 ms to 7,148 ms for neurotypical adults (highlighted by red rectangles in the second row of Panel a). The bar charts (Panel c) compare power between two conditions and two groups during corresponding time windows and at selected channels.

In the mu band (8–13 Hz), autistic adults exhibited sustained significant classification between 6,029 ms and 7,197 ms (*p* clusters < .05, *M*ean AUC = 0.54, Figure 20d), particularly at most the entire brain (*p* clusters < .05, *M*ean AUC = 0.54, Figure 20e). Neurotypical adults exhibited sustained significance between 6,225 ms and 7,197 ms (*p* clusters < .05, *M*ean AUC = 0.55, Figure 20m), particularly in the centroparietal regions (*p* clusters < .05, *M*ean AUC = 0.54, Figure 20n).

During the period for two groups: 6,029 ms to 7,197 ms for autistic adults and 6,225 ms to 7,197 ms for neurotypical adults, we compared theta power at common channels with significant contributions: FP1, FPZ, FP2, FC1, FCZ, T7, C5, C3, CZ, C2, C4, CPZ, CP2, CP4, CP6, P7, P5, P3, P1, PZ, P2, P4, P8, PO7, PO5, PO3, POZ, PO4, PO6, and PO8 (Figure 22). Neurotypical adults (*M* = −3.34, *SD* = 2.33, β = −1.11, *SE* = 0.42, *t* = −2.68, *p* = .009, 95%CI = [-1.94, −0.29]) induced significantly larger suppression than autistic adults (*M* = −2.23, *SD* = 2.01). The incongruent condition under no-pain events (BG-FE+; *M* = −2.97, *SD* = 2.10, β = −0.29, *SE* = 0.06, *t* = −3.37, *p* < .001, 95%CI = [-0.41, −0.16]) elicited significantly larger mu suppression than the incongruent condition under pain events (BG+FE-; *M* = −2.68, *SD* = 2.40). No significant interaction effect was found (*p* > .05).

**Figure 22.**
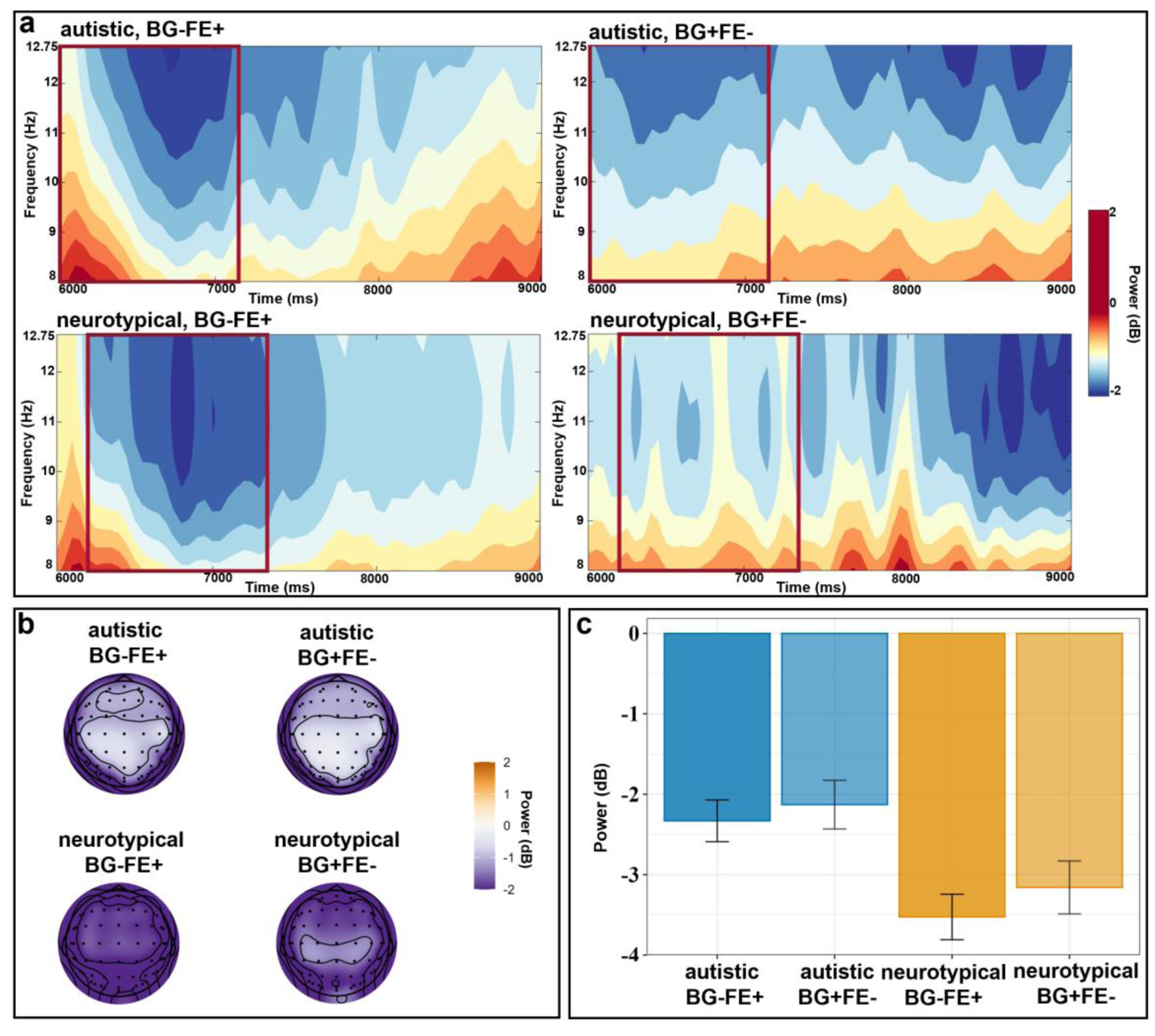
ERSP Comparing BG-FE+ and BG+FE-During the Facial Expression Phase in the Mu Band. *Note*. Panel a: ERSP power distributions across two conditions for two groups over time from the frontal to parieto-occipital channels. The topographic maps (Panel b) illustrate power distribution from 6,029 ms to 7,197 ms for autistic adults (highlighted by red rectangles in the first row of Panel a), and from 6,225 ms to 7,197 ms for neurotypical adults (highlighted by red rectangles in the second row of Panel a). The bar chart (Panel c) compares power between two conditions and two groups during corresponding time windows and at selected channels.

## Discussion

The present study has addressed a critical gap: how the congruence between different pain expressions influences the neural mechanisms of pain empathy in both autistic and neurotypical adults. In addition to the pain empathy paradigm established in Wang et al. (2025), we manipulated the congruence (congruent versus incongruent) between these pain expressions under both pain and no-pain conditions. Furthermore, we employed both multivariate and univariate analyses to systematically investigate temporal dynamics and neural oscillatory activity during different pain response phases. In sum, both autistic and neurotypical adults engaged the N2-P3 ERP complex, as well as theta and mu activity, although the degree of neural engagement varied between groups and across conditions. During the body gesture phase, neurotypical adults exhibited distinct neural representations between the pain and no-pain conditions, which was also evident in a negative ERP pattern. However, autsitic adults showed no significant between-condition differences. During the facial expression phase, we observed autism-specific neural patterns that suggest difficulties in both empathy and conflict resolution. The following section discussed these key findings and their implications.

### Distinct Neural Mechanism for Empathy and Conflict Resolution in Both Groups

We observed distinct neural pathways underlying conflict resolution and empathy, which depended on the congruence between pain expressions. Both processes were observed in autistic and neurotypical adults, suggesting that both groups share neural architectures for processing naturalistic social cues (Hasson et al., 2009). Specifically, when individuals viewed a neutral body gesture (BG-), the following incongruent facial expressions with pain (BG-FE+) elicited a distinct neural representation compared to congruent faces (BG-FE-). This difference merged as early as approximately 100 ms after the onset of the facial expression phase, as shown by multivariate pattern analysis (MVPA). Subsequent univariate analysis identified this difference as a heightened centroparietal N2 for the incongruent pairing (BG-FE+). The increased N2 amplitude indicates both an attention allocation to emotional faces (Liu et al., 2018) and a swift detection of the error (Chmielewski & Beste, 2017). We speculate that individuals allocate more attention to the mismatch between the expected no-pain faces and the unexpected painful faces.

In the late stage (∼400 ms after the onset), the incongruent faces (BG-FE+) elicited a larger centroparietal P3 than the congruent faces (BG-FE-). This implies enhanced cognitive reappraisal to resolve the conflict between expectation and reality (Chmielewski & Beste, 2017; Liu et al., 2018). While two prior ERPs studies in neurotypical adults observed that early ERPs were sensitive to static body-face incongruence (Meeren et al., 2005; Li, 2021), our dynamic and naturalistic pain paradigm revealed a dual-stage process of conflict detection and resolution.

MVPA and ERSP analyses identified a theta power increase (176 ms after the onset) for the incongruent pairing (BG-FE+) compared to the congruent pairing (BG-FE-). While prior studies suggested that early conflict monitoring is reflected in frontal theta power enhancement (Muralidharan et al., 2023; Xie et al., 2025), we observed a more widespread theta distribution extending from the frontal to the occipital regions, which are involved in memory encoding and retrieval (Herweg et al., 2020). The co-occurrence of the N2-P3 complex and theta activity aligns with established neural biomarkers of cognitive control (Harper et al., 2014; Wienke et al., 2018), but the widespread theta activation suggests an additional role for the memory system within a predictive coding framework (Friston, 2005), This is particularly relevant in a dynamic social processing, where individuals continuously generate and update prediction errors by recalling prior stimuli and encoding incoming social inputs (Sinclair et al., 2021).

Under the pain condition where protagonists responded with painful body gestures, the congruent pairing (BG+FE+) increased the N2-P3 complex and theta activations compared to the incongruent pairing (BG+FE-) in both groups. MVPA analysis enabled us to differentiate the role of the N2-P3 complex in empathy from its role in conflict resolution, based on different time windows and spatial distributions. Because bottom-up affective arousal often occurs earlier than top-down error detection (McRae et al., 2012; Padilla et al., 2014), the N2 in empathy appeared earlier than the N2 in conflict resolution. Consistently, the empathy-associated N2 was larger in the most frontal region of the scalp, which is linked to emotional arousal (e.g., Fan et al., 2014; Vecchio & De Pascalis, 2023). Specifically, the enhanced early frontocentral N2 (around the onset of stimuli) and the late parietal P3 (200 ms after the onset) partially aligned with the temporal dynamics of empathy: an early emotional arousal followed by a later cognitive evaluation of pain cues (e.g., Sun et al., 2017; Vecchio & De Pascalis, 2023). Our findings further indicate that this empathy hierarchy exists in autistic adults, and highlight the role of pain context in shaping facial expression perception. Congruent prior body gestures enhanced the perceived genuineness of facial expressions, thereby facilitating an empathic concern (Hadjistavropoulos et al., 2011; Hill & Craig, 2002).

Simultaneously, we observed a theta-mu pattern with a larger theta increase and a larger mu suppression in the congruent condition (BG+FE+) than the incongruent condition (BG+FE-) in both groups, indicating increased emotional sharing (theta; Lavín et al., 2023; mu et al., 2008) and pain resonance (mu, Jelsone-Swain et al., 2023; Yang et al., 2009) to congruent pain cues. Although mu suppression was absent when comparing BG+FE+ with BG-FE-in the current study, the significant mu suppression in the BG+FE+ versus BG+FE-comparison highlights how the congruence influences the mirror neuron system (MNS) activation during pain observation. Expected actions primarily activate the MNS, while unexpected actions recruit the mentalizing system to resolve prediction errors before mimicry begins (Mou et al., 2024; Raz et al., 2020). Thus, when facial expressions were congruent with body gestures (BG+FE+ or BG-FE-), the MNS engagement was enhanced in both conditions, which thereby reduced the between-condition differences. In contrast, the incongruent condition BG+FE-demonstrated an unexpected facial expression, which appeared to disrupt the MNS activation, thereby enlarging the between-condition differences. Despite ongoing debate regarding mu suppression in pain empathy (Hobson & Bishop, 2017), our findings imply that mu suppression is not modulated by pain cues alone, but rather by the congruence between multiple pain expressions.

### Autistic Adults Exhibited Challenges in Empathy and Conflict Resolution

Despite shared neural architectures for empathy and conflict resolution, we observed quantitative differences in neural markers between autsitic and neurotypical groups, suggesting an altered dual-process neural activation in autistic adults. First, MVPA analysis for the body gesture phase revealed that neurotypical adults showed an early neural sensitivity to pain expressed via body gestures (∼200 ms after the onset of body gestures). This sensitivity was identified by the increased N2 amplitudes for body gestures with pain (BG+) compared to no-pain ones (BG-), indicating greater affective arousal to others’ pain in neurotypical adults (e.g., Fabi & Leuthold, 2017, Jiang et al., 2020). However, autistic adults showed no between-condition difference, inconsistent with one prior study that observed heightened emotional arousal to pain in autistic adults (Fan et al., 2014). This discrepancy could be due to the type of pain stimuli. While images of painful body limbs are often exaggerated and explicit, potentially triggering heightened alertness in autistic individuals (Bogdanova et al., 2022), our dynamic displays of body gestures represented a more naturalistic avoidance of pain, which requires both biological motion perception and the understanding of motion intentions that autistic adults are known to have difficulties in processing both (Boria et al., 2009; Freitag et al., 2008).

In the congruent pain condition (BG+FE+), autistic adults showed a larger emotional arousal than neurotypical adults. Furthermore, the difference in emotional arousal between the congruent (BG+FE+) and the incongruent (BG+FE-) conditions was greater in autistic adults than in neurotypical adults. Although larger early ERPs to faces have been suggested to reflect greater social approach in autism (Jones et al., 2018), we interpreted this result instead as an altered alertness when perceiving others’ pain due to hypersensitivity to sensory input in autism (Marco et al., 2011). This atypical early ERP pattern was coupled with altered mu suppression. Specifically, during the late stage, autistic adults exhibited a smaller and slower (classification significance onset was 200 ms after the one for neurotypical adults) mu suppression difference between the BG+FE+ and BG+FE-conditions compared to neurotypical adults. The slower mu suppression, which is a signature neural pattern linked to autism traits (Strang et al., 2022), is specifically associated with their impaired empathy. In contrast the comparable mu suppression reported by Fan et al. (2014) when autistic and neurotypical adults viewed others’ painful body limbs, our finding highlights the influence of stimulus dynamics on mu activation patterns. Mixed findings on mu suppression across autism studies have been attributed to differences between static and dynamic stimuli (Hamilton, 2013), or between social and non-social stimuli (Chan & Han, 2020). The present study implies that mirror neuron system dysfunction in autism (Oberman et al., 2005) is more predominant in dynamic and multimodal pain scenarios. This attenuated mirror neuron system activation may underlie diminished empathic concern (Fecteau et al., 2006; Genzer et al., 2022) which was reflected in the lower ratings of empathic concern given by autistic adults compared to neurotypical peers.

The pain comparisons under congruent conditions (i.e., BG+FE+ versus BG-FE-) revealed a larger P3 difference in autistic adults than neurotypical peers. This result aligns with prior studies that observed an enhanced P3 response to pain stimuli in autistic adults (Fan et al., 2014; Li et al., 2022a), suggesting that autsitic adults allocate excessive cognitive resources to reappraise others’ pain. However, autistic adults did not show a significantly larger P3 when comparing the BG+FE+ and BG+FE-, indicating that additional cognitive resources were consumed to resolve the errors caused by the incongruence between painful body gestures and neutral facial expressions. The enlarged P3 for pain evaluation in the BG+FE+ and the enlarged P3 for conflict resolution in the BG+FE-led to a reduced P3 difference between conditions.

When comparing two incongruent conditions (BG-FE+ versus BG+FE-), autistic adults exhibited significant differences in early-stage ERP, showing a larger centroparietal N2 for BG-FE+. During the late stage, autistic adults exhibited a significantly larger but slower centroparietal P3 for BG-FE+ than for BG+FE-, with a larger condition difference than neurotypical adults. The enlarged N2-P3 complex amplitudes suggest not only an increased early-stage sensitivity to conflict errors but also greater late-stage cognitive efforts to resolve these conflicts (Harper et al., 2014; Wienke et al., 2018). While both groups engaged in conflict resolution for BG-FE+, only autistic adults showed quantitatively greater engagement. This selective sensitivity to BG-FE+ suggests a specific difficulty in processing conflicts related to salient facial expressions in autistic adults. Compared to neutral faces, facial expressions with pain serve as explicit and reliable signals of danger (Kappesser, 2019). Their pairings with neutral body gestures might create an “unrealistic” prediction error contradicting genuine pain situations (Enzi et al., 2016; Zanelli et al., 2025). Neurotypical adults may flexibly adjust their expectations and downregulate these prediction errors (Sapey-Triomphe et al., 2023). However, due to reduced flexibility in re-evaluating prediction errors in social contexts (Lage et al., 2024; Van de Cruys et al., 2014), autistic adults may activate a persistent and overwhelming conflict resolution process.

Beyond these condition-modulation effects, we observed a consistently larger P3 in autistic adults compared to neurotypical adults across all conditions. While one meta-analysis of neurotypical adults reported that the late-stage ERP is a more robust indicator of empathy than early-stage ERPs (Coll, 2018), we suggest that this persistent and increased late-stage P3 is a strong indicator of autistic-specific social-cognitive processing pattern. This interpretation is based on the established link between P3 amplitudes and individual differences in executive function (EF; Reed et al., 2022). Autistic individuals have shown EF impairments, particularly in highly realistic tasks (Kenworthy et al., 2008; Demetriou et al., 2018). Therefore, we speculate that our current paradigm, which integrated pain perception and conflict resolution, placed an excessive burden on the EF resources of autistic adults. The increased P3 reflects their overloaded cognitive effort in implementing top-down cognitive control when facing complex and naturalistic social scenarios.

### Implications and Limitations

Despite the novel pain empathy paradigm and advanced analytical approaches, several limitations should be considered in future research. First, although this study was a pioneering EEG investigation on this topic in autistic adults, we did not examine potential adult-specific confounding factors. Unlike children, adults occupy multiple social roles in family, work, and broader social communities, where socioeconomic disparities can influence prosocial behaviours (Jiménez-Moya et al., 2021). Additionally, adults gain more social experiences over time, such as increased exposure to traumatic events that can shape their perception of others’ pain (Tesarz et al., 2020). Therefore, future studies on autistic adults should incorporate broader assessments of personal information, such as socioeconomic status and trauma history. Second, although using animations to depict dynamic pain events is more naturalistic than showing static pictures, the absence of real human actors, such as ones in live-action videos, may still limit the ecological validity. Future research could involve trained actors to perform pain, though such approach may involve greater expense and require a high level of performance fidelity.

This study has provided both theoretical and practical implications. First, this study advances the current understanding of pain empathy in autism by pioneering the examination of pain expression congruence based on the predictive coding framework. While prior studies have linked atypical prediction coding in autism to their core symptoms, such as hypersensitivity/hyposensitivity, social interaction difficulties, and repetitive behaviours (Arthur et al., 2023; van Boxtel & Lu, 2013), this is the first study to apply this framework to examine pain empathy in autism, where current research yields inconsistent findings (Shalev et al., 2022). Future studies should explore the challenges when autistic individuals face with uncertainty prediction in social contexts. Additionally, this study highlights the influence of pain context on empathic processing in autistic adults. Empathy is a multifaceted capability with different subcomponents that are activated through both bottom-up and top-down processes shaped by context (Heyers et al., 2025). Previous pain empathy studies on autism have largely relied on isolated pain stimuli (i.e., body limbs, facial expressions, or body gestures), neglecting the dynamic and multimodal nature of real-life pain expressions. The present study revealed specific difficulties in top-down processing of a more naturalistic and complex pain context for autistic adults. Future autism research should prioritize ecologically valid paradigms that are distinct from controlled laboratory settings to test real-life challenges for autism.

To our knowledge, this study is one of the few to investigate the neural mechanisms underlying social processing in autistic adults, a group receives less research and clinical attention than autistic children. In reality, autistic adults face broader challenges, such as securing a job, or maintaining social relationships, and managing independent livings. The present study challenges the common assumption that aging could develop compensatory mechanisms, instead highlighting the great difficulties autistic adults have in understanding others’ pain. The overwhelming cognitive effort they expend would cause their failure in maintaining social interactions and developing mental illnesses. We suggest more clinical interventions specialized for autistic adults, focusing on training their social skills for complex real-life social contexts, such as workplace, home and public places. Additionally, researchers should recruit autistic individuals from different age ranges to investigate any age-related developmental trajectories underlying social processing in autism.

## Conclusion

The current study developed a pain empathy paradigm to examine how the pain expression congruence influences the neural mechanisms underlying empathy in autistic and neurotypical adults. The integration of multivariate and univariate pattern analyses dissociated two distinct cognitive processes, i.e., conflict resolution and empathy, under different pain congruence conditions. These neural activities were reflected in the N2-P3 ERP complex, as well as theta and mu neural bands. Compared to neurotypical adults, autistic adults exhibited difficulties in both processes, but with different neural patterns. Specifically, autistic adults exhibited heightened automatic emotional arousal but weaker emotional resonance and expended more cognitive efforts to evaluate others’ pain situations. When faced with others’ painful facial expressions preceded by an incongruent body gesture, autistic adults consumed more cognitive resources to resolve the conflict. The consistently larger P3 in autistic adults across all conditions suggests that they encounter persisting difficulties in processing social stimuli.

## Acknowledgments

This project has been supported by the Hong Kong Research Grants Council (RGC)-General Research Fund (17620520) and Research Fellow Scheme (RFS 2021-7H05).

